# Modelling symplasmic transport: from biophysical basics to steeper root gradients

**DOI:** 10.1101/2025.01.01.631002

**Authors:** Eva E. Deinum, Athanasius F. M. Marée, Yoselin Benitez-Alfonso, Verônica A. Grieneisen

## Abstract

Development and spatial pattern formation are inherently linked. The coordinated determination of cell fates is crucial in any developmental process and requires extensive intercellular communication. Plants cells exchange many molecular signals via the symplasmic pathway, i.e., via plasmodesmata: narrow channels connecting the cytoplasm of neighbouring cells. Regulation of symplasmic transport is vital for normal plant development, and mutations that disrupt this regulation are often embryo or seedling lethal. In many tissues, symplasmic transport of small molecules is diffusion driven, resulting in a non-selective and bidirectional transport, although net directionality could arise from gradients. This has led to the (dogmatic) belief that symplasmic transport can only be detrimental to pattern formation, because signalling molecules cannot be confined, and gradients would fade. Here, we develop a detailed biophysical description of symplasmic transport to explore how plasmodesmata affect gradients in a linear tissue. We then apply the model in more complex tissue contexts, observing and explaining, e.g., that symplasmic transport may result in *steeper* gradients in the root apical meristem. In conclusion, our model provides a reference framework for estimating the consequences of symplasmic transport and explains how symplasmic transport can contribute to more robust developmental patterning.

## 1 Introduction

Development is not only a temporal, but also an inherently spatial process. The process involves both coordination of developmental decisions among cells and cell types and, subsequently, the isolation of these decisions to individual cell (type)s. Owing to the thick cell walls that surround and immobilize their cells, plants have evolved a unique set of signaling mechanisms for this coordination. To understand how specific signaling molecules can form informative gradients and/or interact to control development, it is essential to understand how they move through tissues.

An important factor is neighbour-to-neighbour intercellular communication via plasmodesmata, narrow channels connecting the cytoplasm of neighbouring cells. The aperture of plasmodesmata is controlled through the deposition and degradation of callose, by callose synthase (CalS a.k.a. GSL gene family) and *β*-1,3-glucanase respectively, with further regulation by other protein factors [81]. Callose deposition is assumed to be a fast process and is also involved in the closure of plasmodesmata in response to wounding [44] and pathogen infection [25]. Callose regulation is essential for plant development and mutants defective in this regulation are often embryo or seedling lethal [36, 5, 78]. Despite the fact that mutations in the regulation of this so-called symplasmic transport are often embryo or seedling lethal, this transport mechanism is often conveniently overlooked in modelling approaches.

This is well illustrated by the story of auxin, a plant hormone that has long dominated the study of plant development (e.g., see [8, 28]). Auxin informs the different developmental zones in the growing root tip through a long range gradient [29, 50]. Auxin gradients also control tropic responses [70], and its (patterned) local accumulation is essential in guiding phyllotaxis [61, 35, 69] and (root) lateral organ initiation [56, 42, 19] – to name a few examples. Most attention in auxin related studies goes to its active transport by the polarly localized PIN proteins and other carriers [23]. Given its small size, however, auxin can also easily move through most plasmodesmata. Symplasmic transport impacts auxin gradients [64, 31, 52, 21]. Callose mediated closure of plasmodesmata is essential for tropic responses, which depend on local auxin gradients [31], as well as leaf vein patterning [47]. Symplasmic transport is also actively regulated during lateral root emergence [66] and can impact the distribution of other plant hormones. Strikingly, root nodule primordium formation (in the indeterminate nodule forming model species *Medicago truncatula*) depends on the degradation of callose in cell walls, a process associated with increased symplasmic transport [22]. An important early signal in nodulation is cytokinin [74], which has recently been reported to increase symplasmic transport in leaves [33]. Plant cells can also send advanced messages to their neighbours in the form of transcription factors and small RNA molecules that move symplasmically [48, 54, 27, 50, 51].

Symplasmic transport can be divided into two types. The first is generic: the passive movement of all sufficiently small molecules. This is called *non-targeted symplasmic transport* and is typically diffusion driven [68, 51]. The second is specific and therefore called *targeted symplasmic transport*.

This is a container term for a variety of different mechanisms that allow symplasmic movement of molecules that, typically, would not be able to pass in absence of the mechanism [80]. Some forms of targeted transport affect subsequent non-targeted transport. In particular, some viral “movement proteins” induce structural alterations of the plasmodesmata, which affect the non-targeted transport properties [49, 6].

The boundary for being “small enough” to pass through plasmodesmata is conceptually referred to as the “size exclusion limit” (SEL) and mostly depends on a molecule’s hydrodynamic dimensions [71, 17]. The SEL is developmentally regulated and varies among different tissues, developmental stages and different cell faces, as does the number and density of plasmodesmata [82, 83, 62, 45, 20].

For development, however, the relevant measure may not be whether a signalling molecule *can* pass to a neighbouring cell, but how efficiently this happens. This is better captured by a measure called the “effective symplasmic wall permeability” for the substance(s) of interest. More recent measurements focus on the functional determination of symplasmic wall permeability based on photobleaching and photoactivation approaches, using mostly fluorescein dyes and GFP-like molecules [46, 64, 55, 24]. One would naively expect that functionally obtained effective permeability values would roughly match those from model calculations based on plasmodesmata counts and geometries observed with electron microscopy. This, however, is not always the case, with so-called type I plasmodesmata (lacking a clear cytoplasmic sleeve for transport) as a notorious example [55, 79]. Resolving this conundrum requires advanced models at both the microscopic [18] and tissue level.

Despite a few modelling studies that include a symplasmic component [50, 52], a systematic biophysical model at the tissue level is currently lacking. Here, we will focus on the non-targeted transport of generic small molecules. At the molecular level, this transport mechanism has no bias for any specific direction. In the presence of a (turgor) pressure gradient, the direction of non-targeted transport could be biased by the pressure induced hydrodynamic flow.

Simulations of root water uptake suggest that this process induces a strong directionality of symplasmic transport over only a few cell-cell interfaces, particularly where apoplasmic water fluxes are blocked [14]. Other very specific contexts include secreting trichomes [12].and the phloem, which also contains highly specialized plasmodesmata [75, 13, 63]. Otherwise, non-targeted transport is considered to be diffusion driven [51]. We, therefore, do not consider hydrodynamic flow in our model.

We start investigating the basic biophysical properties of non-targeted symplasmic transport by constructing and studying gradients in a 1D model. In this model, we consider both the cellular compartmentalization of the tissue and intracellular diffusion of the signal (Figure 1). We show that although intracellular diffusion is often ignored in plant models (including most models of leaves and the shoot apical meristem [39], and even some recent root models [52]), assuming a homogeneous intracellular concentration can result in a drastic overestimation of a signal’s mobility and range. This simple model setup allows us to cross-validate analytical and simulation results, derive the scaling behaviour of non-targeted symplasmic transport and contrast this with apoplasmic transport. These results could be directly applied in studying symplasmic transport in moss protonema, because these strings of cells strongly resemble a 1D tissue. We subsequently considered our models in 2D to investigate the effects of tissue anisotropies, which are a natural feature of almost all plant tissues. Finally we investigate how the regulation, or, in mutants, lack thereof, of symplasmic transport can affect root gradients. Together, our findings provide a reference framework to assess the impact of symplasmic transport under various conditions.

**Figure 1:**
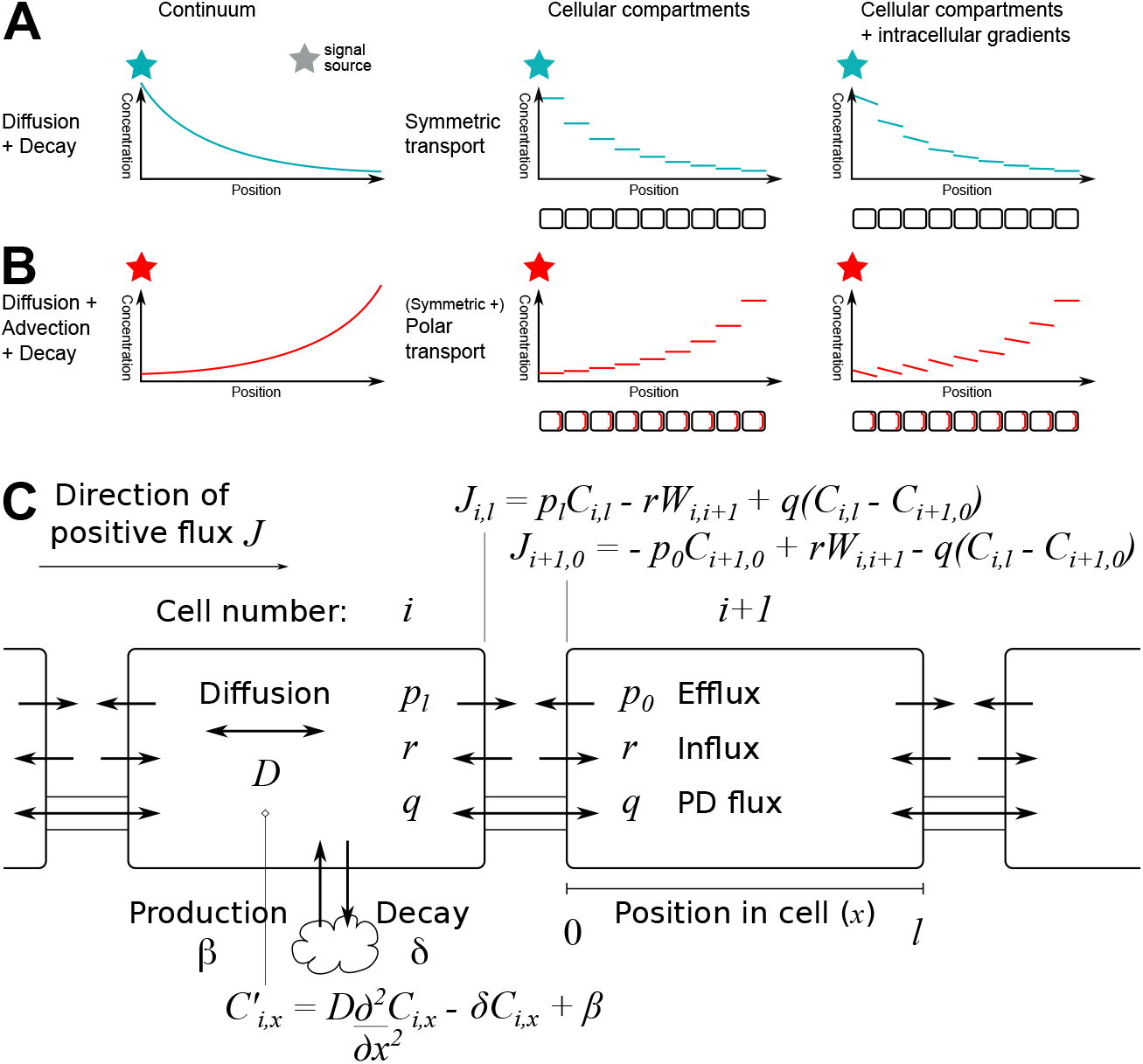
Morphogen gradients could span many cells, and are affected by cellular compartmentalization as well as intracellular gradients. **A**: Pure symplasmic transport resembles diffusion + decay in its possibilities. A diffusion decay model is a continuum model. A more precise model considers that the tissue is made of individual cells/compartments. Inside cells, particularly is they are long, intracellular gradients arise. Colored stars indicate the source of the signal. **B** The combination of symplasmic and directed apoplasmic transport resembles diffusion + advection + decay. At a cellular level, this is driven by polar transport, as indicated by the red accents (export proteins) in the cell file cartoons below the cellular models. **C**: **Model outline with equations**. Our model considers both cellular compartments and intracellular gradients, from which a tissue level profile naturally arises. All fluxes through the walls are modelled using effective permeabilities (units *µm/s*). Concentrations are given as *C*_*i,x*_ in cell *i* at location *x* and *W*_*i,i*+1_ in the wall between cell *i* and *i* + 1. Model parameters: decay constant *δ*, cell length *l*, effective wall permeability *q* and diffusion coefficient *D*.

## 2 Results

### 2.1 Model definition and sketch of solutions

The plasmodesmata allow for diffusion of sufficiently small particles, which move via the cytoplasmic sleeve (symplasmic space between the plasma membrane and the desmotubule). This process can be captured as an effective diffusive permeability *q* for each interface separating two cells. The value of *q* will depend on the substance, with a very strong influence of the particle’s hydrodynamic dimensions, and the plasmodesmata themselves [71, 17, 18]. For the small (0.5 kDa) molecule fluorescein, measured values are typically in the range of 1-10 *µms*^−1^ [26, 64, 46].

For simplicity, we first define our model in 1D (Figure 1C). Here, the effective permeability *q* results in a flux *J*_*i,i*+1_ = *q*(*C*_*i,l*_ − *C*_*i*+1,0_) over the wall between consecutive cells *i* and *i* + 1, with *C*_*i,l*_ the concentration at the far end of cell *i* and *C*_*i*+1,0_ the concentration at the near end of cell *i* + 1. Within each cell the substance moves by diffusion (with diffusion coefficient *D*), with decay (with rate *δ*) occurring everywhere in the cell. Production (with rate *β* per volume) may take place in designated cells.

To see how far a signal could spread, we first solved for the steady state of the model starting from the general steady state concentration profile in single cell, which contains only diffusion and decay. We then chained together consecutive cells using the fluxes over the wall that separates them. For details, see appendix A.2. We cross-validated our results with numerical simulations of a long line of cells with production in the middle cell only and parameters reasonable for a small molecule like fluorescein or a plant hormone (Figure S1).

In a coarse-grained form, i.e., a continuous function that matches the full solution at a single position per cell, the steady state resembles the solution of a simple diffusion-decay process and has the same mathematical structure. We exploited this structure to introduce a concept of a characteristic length (see below) and derive an analytical approximation for the dynamics of the system (appendix A.3), which was in good agreement with the numerical simulations (Figure S1).

### 2.2 Spread of a signal using characteristic lengths

To investigate how the range of the signal is affected by different parameters, we used the concept of a characteristic length, here defined as the distance over which the (steady state) concentration declines by a factor 1*/*e≈ 0.37 (illustrated in figure 2A). A characteristic length is, for example, often used to describe morphogen gradients [76].

**Figure 2:**
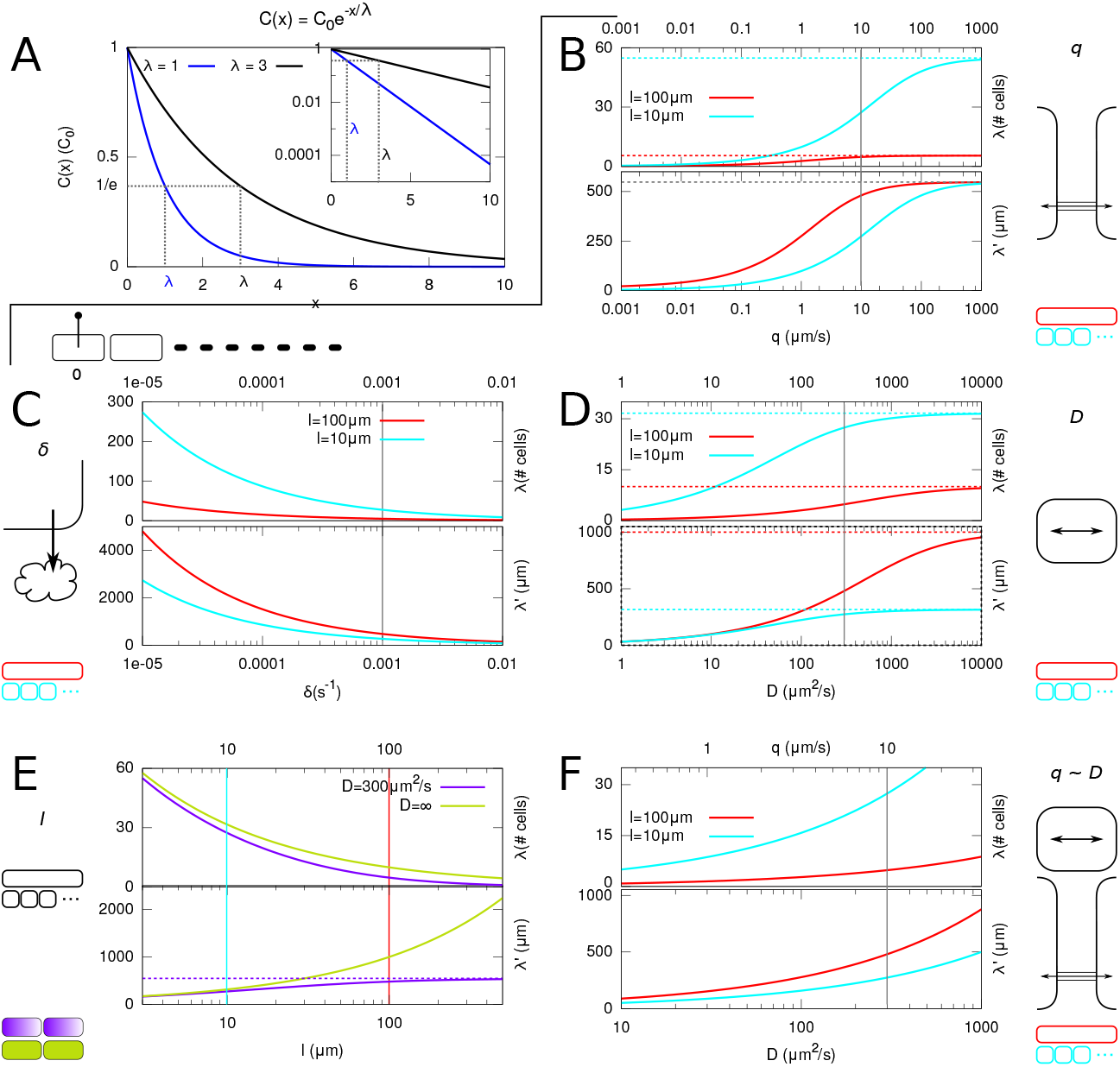
Characteristic lenghts: Comparing and contrasting morphogen gradients. Here we will access the impact of each independent parameter to the steady state morphogen gradient. **A**: The steepness of the concentration profiles is given by itscharacteristic length, the length over which the concentration declines by a factor 1*/e* (≈ 0.37); the blue morphogen gradient declines more quickly than the black gradient, and indeed, the the blue lambda (1) is smaller than the black gradient (3). This measure can be expressed in number of cells (*λ*) or physical length (*µm*; *λ*^*′*^). **B-F**: Dependence of *λ* (upper panel) and *λ*^*′*^ (lower panel) on individual parameters (B: permeability *q*, C: decay rate *δ*, D: diffusion coefficient *D*, E: cell length *l*), as indicated with symbols and cartoons at the sides, for two cell lengths (**B**,**D-F**) or with (purple) or without (light green) intracellular gradients (**E**). Default parameters: *q* = 10 *µm/s, δ* = 0.001*s*^−1^, *l* = 100 *µm, D* = 300 *µm*^2^*/s*. As both *q* and *D* depend on particle size, we also calculated the characteristic lengths keeping the ratio *q/D* fixed (**F**).

It is not *a priori* clear what is more limiting to the spread of a signal: the physical distance it has to travel, or the number of walls it has to cross. We, therefore, computed the characteristic length in two different ways, denoted *λ* for number of cells, or *λ*^*′*^ for physical distance (in *µm*). Figure 2B-E shows the effect of each model parameter on the characteristic lengths for two different cell lengths per graph.

Obviously, the characteristic length increases with increasing wall permeability *q*, but this is limited by intracellular diffusion (Figure 2B). The characteristic length also increases if the signal is more stable (lower *δ*) (Figure 2C). Here it is important to note, however, that *δ* also sets the time scales of reaching the steady state (Figure S1B-D), as time always appears in a factor 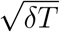 in the temporal solution (Equation 69, appendix A.3). With increasing diffusion coefficient *D*, the characteristic length increases, this time in a permeability and degradation limited fashion (Figure 2D). The latter explains why the signal spans more cells with short cells, but more distance with long cells.

In none of these graphs, the two curves for different cell sizes overlayed, indicating that both cellular scale and physical distance are important for the range of the signal. For cell length *l*, an even more extreme picture emerged: with increasing *l*, the average number of cells travelled decreases, but the distance increases (Figure 2E). Even with infinitely fast intracellular diffusion, *λ* decreased with increasing *l*, because the signal is slowed down at each wall (barrier) and the total degradation per cell is proportional to its volume. Notably, explicitly considering intracellular diffusion reduced *λ* by 13% for the high default value of *D* = 300 *µm*^2^*/s* [29, 42] and short cells (*l* = 10 *µm*) and by even 52% for long cells (*l* = 100 *µm*). Only in the case of effective wall permeability *q* (fig. 2B), the long and short cells share the same upper limit for the characteristic length in physical distance 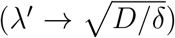. In this limit, the walls no longer pose any barrier, and the system reduces to diffusion/decay in a homogeneous medium.

There is no graph for production rate *β*, as this has no impact whatsoever on the characteristic length. In fact, the whole solution is linear in *β*, because production is not regulated here, so the amount of signal at any point in space and time linearly depends on *β*.

### 2.3 A simplified model captures most model behaviour, but not in extreme cases

Further exploiting the similarity to a plain diffusion-decay process, we defined an effective diffusion coefficient (*δ*-dependent) *D*_*eff*_ = *λ*^2^*δ* (equation 55, appendix A.2.2). We substituted this expression into the time dependent solution of the heat equation. The resulting equation (70, appendix A.3), showed that the short term dynamics is governed by this *D*_*eff*_. This is also visible in figure S1B-D, in which all panels, all with the same *D*, show similar profiles until they approach their respective steady state (where 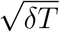 starts dominating).

Taking the limit of *δ* ↓ 0 of *D*_*eff*_, we arrived at a much simpler expression: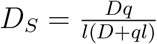 with distance expressed in cell length. Converted to physical distance this is: 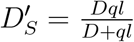 (equations 57 and 58, respectively, appendix A.2.2), a well known result in the context of morphogen gradients in embryogenesis [15] and used in the interpretation of tissue level measurements of symplasmic transport [64]. Both simple and full effective diffusion coefficients gave near identical predictions for a wide range of parameters. The slight underestimation at the shortest time scales appeared to be the result of using a point source rather than a full length producing cell (Figure S2). To find out where the approximation would break down, we increased *δ* to extreme values. In those cases, particularly with low *q* = 0.1 *µm/s*, the full expression for *D*_*eff*_ was required to obtain the correct steady state profile (Figures S1F-G, S3, S4) and both methods underestimated the spread of the signal at the shortest time scales. This underestimation cannot be explained by the point-source, because the difference is more than one cell length. This implies that with high *δ* and low *q*, the dynamics of the coarse grained system is superdiffusive, whereas the steady state can be described as a diffusion/degradation process. In colloquial terms, this can be understood as an extra “effective pulling force” arising from concentration differences over the hard-to-pass walls. Note, however, that this regime is more likely to be relevant for targeted than untargeted symplasmic transport.

The formulas of *D*_*S*_ and *D*^′^_*S*_ elegantly explain that the characteristic lengths cannot be increased indefinitely by increasing the particle’s diffusion coefficient *D* (employing smaller molecules; Figure 2E) or the effective permeability *q* (increasing PD density and/or aperture; Figure 2B) in isolation. Although plants can regulate *q* independent of *D, q* itself linearly depends on *D* [18]. We, therefore, also varied them together with a fixed ratio 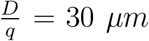 (based on measurements in [64]) which we assume is reasonable for particle sizes much below the SEL (Figure 2F). In this case, mathematical limits are replaced by physical bounds on *D* and *q* like the diffusion coefficient of water in water (≈ 2300 *µm*^2^*/s* [41]).

### 2.4 Comparing symplasmic vs. apoplasmic transport

We then included apoplasmic transport in the same model context as above to address two questions: (i) to what extent can (symmetrical) apoplasmic transport behave the same as symplasmic transport and (ii) how do symplasmic and apoplasmic transport interact?

Assuming that the apoplasmic transport system operates far from saturation, it can be modelled using effective permeabilities, possibly different for influx (*r*) and efflux (*p*) and for different faces of the cell (Figure 1C) [29]. These effective permeabilities capture both passive membrane permeability and carriers mediated fluxes. Using the same method as before, we calculated the coarse-grained 1D analytical steady state profile (appendix A.4).

The influx permeability *r* does not occur in this solution, implying that in 1D the influx capacity does not affect the intracellular steady state concentrations, provided that *r >* 0 is the same for both sides of the wall. The calculations predict that symmetrical apoplasmic transport (i.e., with effective efflux permeabilities *p*_0_ = *p*_*l*_ = *p*) and symplasmic transport yield the same intracellular profiles along lines with *y* = 2*p* + *q* (Figure S5B). Both predictions were confirmed by simulations (Figure S5A). In 2D, this identity breaks down (Figure S5C-F), because the apoplast now forms a continuous network through which it is possible to bypass cells (Figure S5G). The differences increased with decreasing influx efficiency (lower *r*), corresponding with longer apoplasmic travel distances before an exported particle would reenter a cell (c.f. [39]). Such apoplasmic travel, impossible with symplasmic transport or in 1D, has the potential to reduce the impact of tissue topology on patterning processes, as we indeed observed (Figure S6). Nonetheless, the patterns that were identical in 1D remained highly similar in 2D (Figure S5C).

### 2.5 Symplasmic transport can robustly tune gradients of a signal that moves primarily by apoplasmic transport

A key differentiating feature of apoplasmic transport is that it can have a strong directionality resulting from different effective efflux permeabilities at different cell faces (in 1D: *p*_0_ ≠ *p*_*l*_). It has been argued that the efficiency of directed apoplasmic transport could be greatly reduced if the signal can move back through plasmodesmata, for example in the case of auxin [64]. This plant hormone is well known for its directed transport and based on its size is likely to move symplasmically as well. Our 1D model addresses this issue for a string of identical cells, and could effectively model the achievable auxin gradients in moss protonema or a string of BY-2 cells.

Mathematically, a directional bias in the apoplasmic transport does not change the formulas describing the steady state. We call the two characteristic lengths corresponding to both parts of the solution *λ*_*source*_ and *λ*_*end*_ (Figure S7A). From a design perspective, little loss of signal on the way to the root tip is desirable, so *λ*_*source*_ should be large. By the same token, the steepness of the gradient towards the root tip (1*/λ*_*end*_) should be intermediate: steep enough for reliable detection, shallow enough to span enough cells. This model without symplasmic transport has been studied extensively before [53], but we include an overview of model behaviour for easy reference (characteristic lengths expressed in the number of cells: Figure S7). With the default value of *p*_*l*_ or the the default ratio *p*_*l*_*/p*_0_ = 20, *λ*_*end*_ was always smaller than 0.4, meaning a very steep gradient, most likely spanning only a few cells. Notably, the ascending part of the gradient vanished nearly completely for the relatively large value of degradation constant *δ* = 0.01*s*^−1^, although this had very little effect on *λ*_*end*_ (Figure S7AB). We, therefore, replaced the characteristic lengths by more informative measures based on the full steady state solution (Figure 3A). We defined *d*(*X*), the “length of the informative gradient” –which is independent of tissue length (see appendix A.4.1)– and *C*_*N*,0_*/C*_*X*_, the relative concentration increase, based on the point *X* where the ascending (*λ*_*end*_) and descending/flat (*λ*_*source*_) parts of the solution meet.

**Figure 3:**
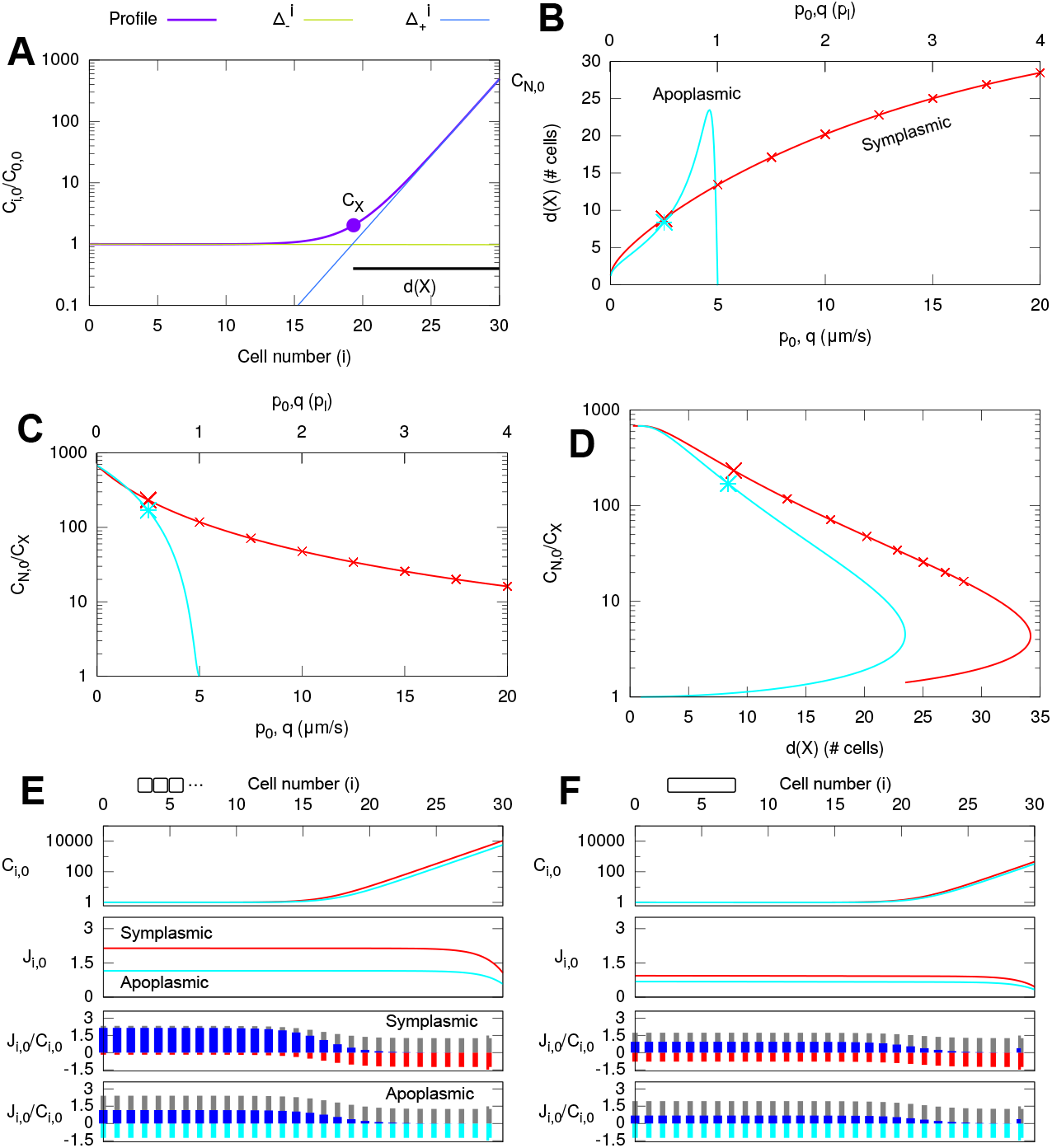
The gradient of directed apoplasmic flux can be tuned more precisely and more favourably with symplasmic than with apoplasmic backflux. Signal is produced in the leftmost cell of the 1D tissue. **A**: Definition of the “length of the informative gradient” (*d*(*X*)) based on the steady state profile and the “relative concentration increase” (*C*_*N*,0_*/C*_*X*_, here with *N* = 30) over this distance (see also appendix A.4.1). **B**: *d*(*X*) as a function of *q* (red: *p*_0_ = 0) and *p*_0_ (cyan; *q* = 0). Markers occur every 2.5*µm/s*. Values of *q* and *p*_0_ are given in absolute values (bottom) and relative to the value of *p*_*l*_ (top). **C**: idem for *C*_*N*,0_*/C*_*X*_. **D**: combined curves including markers from B,C.**F**: steady state profiles for symplasmic (red) and apoplasmic (cyan) backflux for parameters as indicated with the large markers in **B-D** (top), corresponding flux through every cell wall (second) and the relative forward (directed apoplasmic, grey) and backward (symplasmic: third, red; apoplasmic: fourth, cyan) and net (blue) fluxes through every cell wall. **E**: idem, but with short cells (*l* = 10 *µm*).

Adding symplasmic transport to the apoplasmic model, we found that allowing for a symplasmic backflux resulted in a less steep, but several fold longer informative gradient (Figure S1H). Based on previous similarities, we wondered to what extent the same could be achieved by reducing the polarity of the apoplasmic transport via increased backflux (*p*_0_). the impact of symplasmic fluxes or apoplasmic backfluxes seemed completely different (Figure 3B,C). Combined, however, the two became a lot more similar (Figure 3D; *q* in the case of symplasmic or *p*_0_ for apoplasmic transport changes along the curve). We consistently found that the symplasmic (*q*) curve was laying outside the apoplasmic backflux (*p*_0_) curve (Figure S8), indicating that a symplasmic flux performs consistently “better” than an apoplasmic backflux in optimizing both *d*(*X*) and *C*_*N*,0_*/C*_*X*_ at the same time. In biological terms this means that with a symplasmic backflux, information could be communicated over a greater distance (across more cells) from the distant end wall and/or it could be more easily detected due to the steeper gradient. The difference was larger with short cells than long cells. The figures also show that tuning the length of a relatively long gradient with apoplasmic backflux requires a very precise control of the backflux *p*_0_, with values only slightly lower than the forward flux *p*_*l*_ (c.f. Grieneisen et al. [28]). With symplasmic transport, on the other hand, tuning can be much more robust. The points at equal *q*-distance in figures 3B-D and S8 show that, depending on the other parameters, it could take unrealistically large values of *q* to complete the *d*(*X*), *C*_*N*,0_*/C*_*X*_-curve.

To better understand why symplasmic backflux always outperformed apoplasmic back-flux, we computed the fluxes over each interface in steady state (appendix A.5.2) for parameters with similar profiles, indicated with the largest symbols on the curves in figure 3B-D. We found that throughout the tissue, the net flux towards the distal end was larger for symplasmic than for apoplasmic backflux (Figure 3E,F). When splitting the flux over each interface in a forward apoplasmic (*p*_*l*_) and backward (either *q* or *p*_0_) part, we found that the normalized backflux by the apoplasmic pathway was the same everywhere, whereas the symplasmic backflux was smaller in the “flat” part of the profile. Only towards the end, symplasmic backflux reached the same level as with apoplasmic backflux. This difference, hence its impact on the net flux, was strongest with small cells (Figure 3E). Similarly, the steady state was also reached faster with symplasmic backflux and the difference in reaching the steady state was larger for the small cells.

### 2.6 Higher dimensions: effects of different anisotropies

2D tissues have more degrees of freedom, which can give rise to anisotropies due to cell aspect ratios, different effective permeability for different faces of the cell [82, 10], and the alignment of cells and cell files (“tissue topology”). How do such tissue anisotropies affect the spreading of a biochemical signal?

PD density counts suggest that cell files of a tissue are coupled, although the longitudinal walls have lower PD densities than the transverse walls in young roots [82]. To investigate the impact of such coupling, we created a long strip of 2D tissue with different effective permeabilities for transport within cell files (*q*_*l*_: for longitudinal transport) and between (*q*_*t*_: for transverse/radial transport) with a single producing cell (Figure 4A-F). We observed how fast the steady state profile through the producing cell (red) decayed from the 1D profile with the same parameters (cyan) to the diluted 1D profile where the signal is equally distributed over all 11 cell files (blue). Comparing all graphs, we found that the value of *q*_*t*_ was more important than the ratio *q*_*l*_ : *q*_*t*_ (Figure S9). This implies that the lateral coupling strength between cell files is the most important factor in the transition from local to global behaviour of a signal spreading in elongated organs.

**Figure 4:**
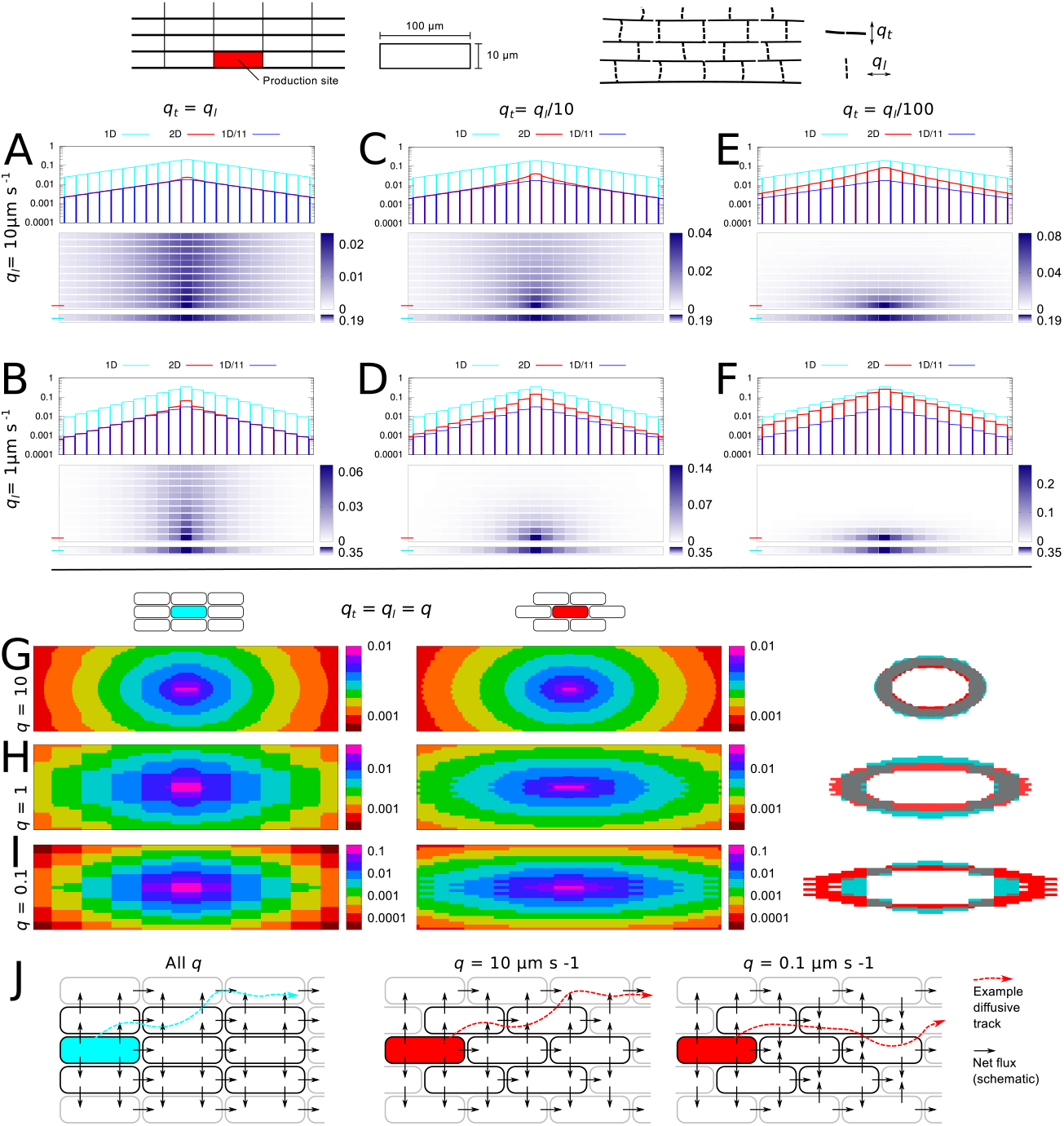
The impact of tissue layout increases with decreasing effective wall permeability. **A-F**: spread of a signal produced in a single cell at the edge of an 11 cell files wide linear tissue with effective permeability *q*_*l*_ within cell files (longitudinal transport) and *q*_*t*_ between cell files. The profile through the producing cell (red) is bound between an isolated 1D profile (cyan) with the same parameters (*q* = *q*_*l*_), and this profile diluted (divided by 11) over all files (blue). **G-J**: Sensitivity to tissue layout depends on *q*. Production occurs in a single cell in the center of the tissue. Concentrations are indicated with different colors using a step function. An overlay of a ring representing the same concentration range for both alignments (cyan: “square”, left; red: “brick”, middle) is shown on the right (gray: overlap). The brick alignment shows a stronger longitudinal transport,particularly in the transport limited regime (**I**). **J**: directions of net symplasmic flux and an example diffusive path. With the brick alignment, net fluxes depend on *q*, resulting in different diffusion dominated (**H**) or permeability dominated (**I**) regimes. Parameters: *D* = 300 *µm*^2^*s*^−1^, cell length: *l* = 100 *µm*, cell width: 10*µm, δ* = 0.001*s*^−1^, *β/*volume = 2*δ, T* = 10*h* (**A-F**) or *T* = 1*h* (**G-I**).

We next investigated the impact of local tissue topology on signal spread, by comparing the two most extreme possibilities for identical rectangular cells in files: a “square” and a “brick” alignment (Figure 4G-I). We used a stepwise color gradient to visualize the concentration profiles around a single producing cell in the center. This resulted in a set of “rings”, each representing a fixed arbitrary concentration range. By overlaying a single equivalent ring of both alignments we found that the brick alignment increased the longitudinal but decreased the transverse range of the signal. With high transport (*q* = 10 *µm/s*), this difference was small, but it became more pronounced with decreasing *q* (Figure 4G-I). The effect can be understood as a shift from a diffusion limited regime to a transport limited regime. In the latter, the number of walls to cross is limiting and the connectivity of the cells becomes important. Only for the brick alignment, the directions of the net symplasmic fluxes per interface differ between diffusion and transport limited regimes (Figure 4J).

### 2.7 Symplasmic transport can increase gradient steepness with the root reflux loop

In the 1D model, the addition of symplasmic transport always reduced gradient steepness towards the distal end of the tissue. It has been shown, however, that this uniform linear setup is not suitable for generating a sufficiently long, but fast establishing gradient as generally assumed in the root [28]. We tested the impact of symplasmic transport in a more realistic root context using the root auxin gradient as an example (Figure 5). We used the same PIN/ auxin efflux permeability distribution and similar cell sizes as in [42]. This root is divided in four zones, starting from the root tip: a three layer thick columella, a”meristematic zone”(MZ) of fifteen 16 *µm* long cells, an”elongation zone”(EZ) of fifteen 60 *µm* long cells, and a “differentiation zone” (DZ) of fifteen 100 *µm* long cells (Figure 5). The PIN distribution in this root gives rise to a reflux loop structure, with rootward transport in the stele and shootward + inward transport in the cortex and epidermis. For symplasmic transport, we split the walls in three groups: H (“horizontal”: radial transport), V (“vertical”: longitudinal transport) and C (“columella”: all walls within the 3 layers of the columella). We compared two cases (both red): high radial (Figure 5A,C) and high longitudinal (Figure 5B,E) symplasmic transport, with a reference case without any symplasmic transport (cyan, figure 5D). Strikingly, the radial symplasmic transport had a much larger impact on the shape and length of the gradient. With high radial symplasmic transport, the auxin depression towards the rootward end of the model “EZ” disappeared. Gradients obtained with combining both high radial and high longitudinal symplasmic transport were similar to high radial transport only, with the exception of a further reduced concentration in the quiescent centre (QC) and columella (Figure S11). The increase in gradient steepness correlated with increased net fluxes from the exterior to the interior cell files in the MZ (Figure 5F-H, S10). Adding high symplasmic transport to the columella did not change these results (Figure S11), possibly because of the high efflux permeability into all directions of these cells.

**Figure 5:**
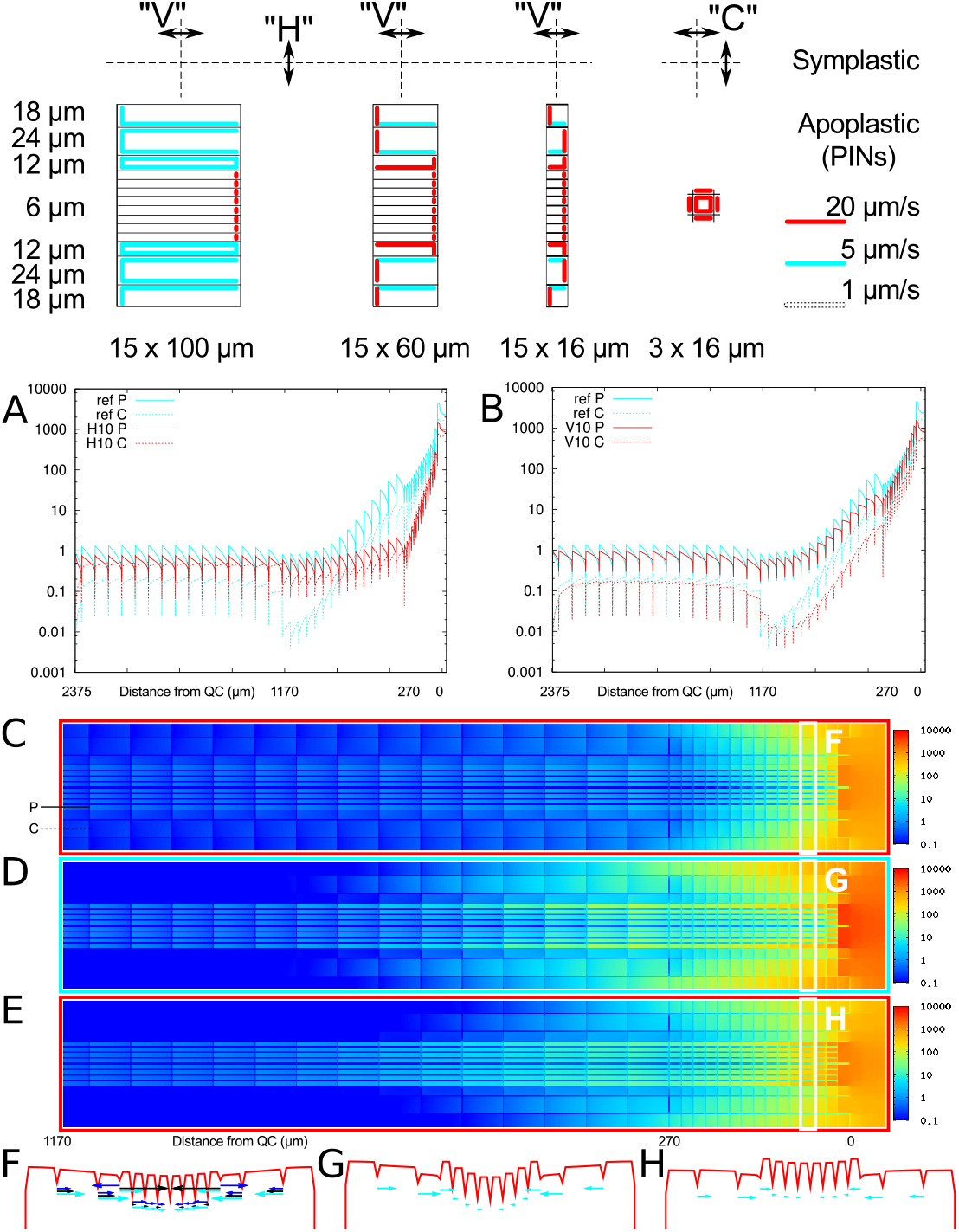
Radial symplasmic transport can create steeper root gradients by enhancing the reflux loop. Symplasmic transport is possible in the radial direction: *H* = 10 *µm/s* (**A,C,F**) or longitudinal direction *V* = 10 *µm/s* (**B,E,H**) of the root (red profiles). Control roots without symplasmic transport (**D,G**) in cyan. **A,B**: Profiles in the pericycle (indicated “P” in **C**) with solid lines, cortex (indicated “C” in **C**) with dashed lines. **F-H** Radial concentration profiles (in red) as indicated by the white squares in **C-E**. Net radial fluxes are indicated by cyan arrows, with arrow length proportional to flux. Where symplasmic transport is possible in the radial direction (**F**), blue arrows show the symplasmic and black arrows the apoplasmic components of the fluxes. Note that the net inward fluxes are larger in **F** than in **G** and **H**. Similar profiles for all (MZ) cell rows in Figure S10. No symplasmic transport in the columella (*C* = 0). *δ* = 5 ·10^−6^*s*^−1^. Roots based on [42], with slightly altered cell sizes. Numbers below graphs and pictures indicate distance from the QC.

## 3 Discussion

We have built a biophysical model of symplasmic transport including intracellular diffusion and explicit apoplastic compartments. From this model in 1D, we have derived effective diffusion coefficients that converge to known results for low substance degradation [15] and yield more accurate results for high degradation and/or low effective wall permeability. When comparing how symplasmic and apoplasmic backflux interact with directed apoplasmic transport [in a string of identical cells], we have found that symplasmic backflux always performs better, i.e., allows for longer and/or steeper gradients for the matched parameters. This is because the symplasmic backflux is depends on concentration differences rather than being a constant proportion of the forward flux. Extending the model to 2D and anisotropic tissues, we have found that symplasmic transport becomes increasingly sensitive to tissue topology and geometry with decreasing effective wall permeability, and this sensitivity is larger than with apoplasmic transport. The latter finding was only possible through explicitly including the apoplast. Finally, we have shown that in more complex contexts symplasmic transport can have counterintuitive effects, like the potential to make the root auxin gradient steeper.

Our model suggests three ways of tuning the range of a signal that travels via symplasmic transport. The first is through diffusion coefficient *D*, which is mostly a direct function of particle size and may be specifically reduced through interactions with other proteins, resulting in larger particles. The second is through effective wall permeability *q*, which again is a function of particle size for non-targeted transport [17, 18], but may be specifically altered through co-factors in targeted transport [80, 49]. Additionally, *q* can be adjusted through callose deposition and degradation [81] as, e.g., in the symplasmic isolation of stomatal guard cells [30]. This process affects *q* for all substances, in a size dependent manner [18]. In line with this, the transcription factors regulating trichoblast/atrichoblast specification in trichome and root hair patterning require different ranges, and the ones acting non-cell autonomously are the smaller members of the set [3, 27]. The third is through degradation rate *δ*, which can be interpreted in a broader sense by including processes that remove a signal from the mobile pool. This happens, e.g., in the SHR/SCR based specification of the endodermis where SCR mediated trapping of mobile, vascular expressed, SHR limits the endodermis to a single cell layer [16]. Conversely, for a long range signal, *δ* has to be low. Indeed, very low values of auxin degradation have been reported for transporting tissues, with particularly low values in pine trees [40]. Modelling studies of the root auxin gradient also use low values for *δ*, ranging from 0.001 *s*^−1^ [52] to 5 ·10^−6^ *s*^−1^ [29, 42]. Even in this range, however, simple 1D models suggest that the chosen value of *δ* could have considerable impact on gradient length (Figure S7A), which appears to be the case when comparing [52] with other studies using lower *δ* [50, 73, 42]. From these considerations on signal range, it follows that long range signals should be small, stable molecules –like all canonical plant hormones. Travel time with diffusive symplasmic transport over longer distances, however, scales quadratically with length, just like simple diffusion. Root-shoot communication and other long distance signaling processes, therefore, relies on mechanisms that scale linearly with length, either by real flow (such as CLE peptides and small RNAs that are transported via the vasculature [57, 43]) or directed (apoplasmic) transport like polar auxin transport [53, 9].

By comparing different tissue layouts, we have observed that the coarse grained spread of a signal is sensitive to tissue topology and cell geometry and that this sensitivity is larger for symplasmic than for similar strength apoplasmic transport and increases with decreasing effective wall permeability (Figure 4, S5, S6). In particular, a more realistic “brick wall” or staggered alignment between cell files tends to increase transport along the cell files and decreases it between files. This could have consequences for modelling studies, which often still use a square alignment of cells, particularly when including intracellular gradients [59, 2, 67, 38, 65, 73]). Similarly, a dramatic impact of tissue topology on the effectiveness long range coordination of cell polarity has been observed in a modelling study on planar polarity factors [1]. Applications that depend on the 1D-derived effective diffusion coefficient like *D*_*eff*_ = *Dql/*(*D* + *ql*) may also be affected by tissue topology, including the quantification of symplasmic transport itself. E.g., the elegant measurements of effective wall permeability for symplasmic transport as in Rutschow et al. [64] involve estimating *D*_*eff*_ from temporarily resolved concentration profiles along the tissue. Based on our results, the impact of these topological effects is expected to depend on cell sizes and decline with increasing permeability. Similar quantification methods for determining effective interface permeability [46] interpret the raw data by considering a single photoactivated cell and its direct neighbour as two homogeneous compartments. Such methods could be enhanced by simulating a larger region of the stimulated tissue.

These considerations on tissue topology may be interpreted as a pledge of basing all model studies on realistic, preferentially experimentally determined, tissue templates. Groups taking this approach, however, tend to model these tissues as a set of coupled ordinary differential equations for computational tractability [4, 52, 77], which ignores intracellular gradients and thus introduces a different type of error. As an illustration, with our default parameters, this would result in a 15% (10 *µ*m cells) or 108% (100 *µ*m cells) overestimation of the characteristic length (*λ*^*′*^) of the gradient for symplasmic transport (Figure 2E). As a golden middle road, models with idealized root geometries can accommodate both intracellular gradients and a realistic overall root shape [72] and even growth [65]. Such models have demonstrated that the narrowing of the root tip and presence of a lateral root cap capture most of the differences between template-based and fully square roots like in Figure 5 [72]. In short, shape matters, but so does growth through the dilution and displacement of signals [2, 65]. The choice of a model formalism, therefore, always involves a compromise. Rutten *et al*. offer a beautiful example of how the impact of geometrical compromises can be compensated and overall realistic behaviour restored through local parameter adjustments, based on a good theoretical understanding from previous models [65]. Our work provides a reference for estimating the impact of different simplifying choices.

Using simplified models along more realistic/complicated ones can greatly increase our mechanistic understanding of the system. Compared to more realistic looking models [52], our square root had the advantage of a straightforward assessment of different symplasmic fluxes. In turn, we can compare our results with the simpler Grieneisen *et al*. model, who found increased gradient steepness by any change that increased the coupling between the exterior (epidermal/cortical) and interior (vascular) layers, increased the steepness of the gradient, including increasing the lateral inward directed PIN efflux carriers in the outer layers and decreasing the number of exterior layers [29]. Radial symplasmic transport similarly increases this coupling, and may need to be suppressed to maintain a sufficiently long gradient. Interestingly, plasmodesmata densities are much lower in the radial than in the transverse walls of the root meristem [82], although effective permeability also depends on the sizes and nature of plasmodesmata, which vary substantially among interfaces [63, 18, 10]. Localized expression of LAX influx carriers in the lateral root cap could further reduce the coupling [34].

The same plasmodesmata densities ([82]) were also used in Mellor et al. [52]. Contrary to our findings, they concluded that symplasmic transport was essential for reproducing a normal auxin gradient (inferred using DII-Venus, which measures auxin activity). A careful comparison of the model parameters shows that, despite very similar shoot-ward:inward auxin efflux ratios, the decoupling between shootward and rootward auxin fluxes in the MZ without symplasmic transport is much stronger in Mellor et al. [52], because of a 16-fold stronger influx capacity in the outer layer (lateral root cap vs epidermis) in their model. Only when they start adding symplasmic transport, which has a radial component, the shootward and rootward fluxes become sufficiently coupled for an effective reflux loop. Once this reflux loop is established, however, both their and our model predict similar effects for comparable changes in symplasmic transport.

In a different root modelling study, Rutten and Ten Tusscher [65] allowed for the symplasmic movement of PLETHORA (PLT) transcription factor proteins, but not auxin. Rutten *et al*. observed, using an idealized full root model with growth, that increased symplasmic PLT transport has a location dependent impact on meristem size, which in the model is determined by a delayed readout of the PLT gradient in the vasculature. An increase of symplasmic transport in the model stem cell niche resulted in a longer meristem, whereas an increase in the adjacent model division zone resulted in a shorter meristem [65]. The first observation can easily be explained in 1D as a consequence of increased PLT loading into the division zone. The second effect seems to result from a flattening of the radial gradients, which reduces the vascular PLT concentration towards the average. We also observed radial flattening with increased radial transport in Figure 5, S10.

These models all consider only a single substance moving symplasmically. In this case, the high level predictions about meristem length with varying radial symplasmic transport are the same, and in line with experimentally observed effects of moderately reduced plasmodesmatal aperture through 35S::GSL8 expression [31]. The underlying mechanisms that the models imply, however, differ tremendously. In other cases, making different choices for which molecules are allowed to move symplasmically may even lead to qualitatively very different predictions. Modification of symplasmic transport also affects other processes involving multiple mobile substances, including stomatal patterning [30, 37], root hair and trichome patterning [27, 3] and lateral root patterning and development [7, 66].

Symplasmic transport is of great importance for normal plant function, but challenging to visualize experimentally and, more importantly, difficult to interpret its importance in relation to other transport mechanisms. This creates a great appreciation of mathematical models in the field. For all of the above examples, and more, the field would benefit from an ecosystem of models. For investigating complex interactions, we need comprehensive multi-level models that combine the effects of symplasmic transport of multiple relevant substances, including hydrodynamic flow where relevant [32], and coherently deriving molecule size dependent transport parameters based on microscopic models considering individual plasmodesmata, like PDinsight and others [18, 58, 11]. Using such microscopic models would be both more realistic and more efficient than exploring transport parameters of the different substances independently. The multi-level models just sketched, however, could easily become computationally expensive and hard to interpret. Simpler models like ours, therefore, play an essential role by providing a biophysical baseline expectation that supports the verification and interpretation of more complex models, and facilitates the functional measurement of symplasmic transport parameters.

## 4 Methods

### 4.1 Overview of mathematical derivations

All mathematical derivations are shown in appendix A. Model equations are shown in Figure 1C and the appendix A. How the steady state profiles are obtained from chaining together intracellular concentration profiles of neighbouring cells is described in appendix A.2 for symplasmic transport only and, using essentially the same method, appendix A.4 for combined symplasmic and apoplasmic transport. In these, the characteristic lengths are defined in appendix A.2.1. Effective diffusion coefficients and their simplified form are defined in appendix A.2.2. The length of an informative gradient for combined symplasmic and apoplasmic transport is calculated in appendix A.4.1. An approximate temporal solution for pure symplasmic transport is derived in appendix A.3. This approach exploits the similarity of the course grained steady state solution to that of the heat equation. The reconstruction of intracellular gradients and local fluxes from the coarse-grained profiles is described in appendices A.5.1 and A.5.2, respectively.

An overview of all model parameters and mathematical symbols used is given in table S1.

### 4.2 Numerical simulations

All numerical simulations are performed on 2D grids, using the same finite volume description of the tissue as in [19]. We use the Alternating Direction Implicit (ADI) algorithm [60], adapted to a band-5 diagonal matrix to guarantee matrix invertibility under all possibly relevant conditions. We found that in cases that could also be solved with a tri-diagonal matrix (e.g., purely symplasmic transport by skipping the wall points), the computational time required for both methods was similar.

The realistic root context (Figure S10, Figure S11) is derived from [42], with minor changes of cell size and the bottom row of vascular cells merged to two QC cells.

## Acknowledgements

The authors thank Bela Mulder and Ton Bisseling for helpful discussions.

The work of EED was funded within the research program of the Netherlands Consortium for Systems Biology (NCSB), which is part of the Netherlands Genomics Initiative (NGI)/the “Nederlandse Organisatie voor Wetenschappelijk Onderzoek” (NWO). Her stay at the John Innes Centre was funded by the European Molecular Biology Organization (EMBO) short term fellowship ASTF 105-2012. YBA work was supported by the United Kingdom Research and Innovation (UKRI) council Future Leaders Fellowship (MR/T04263X/1).

## Author Contribution

EED, YBA and VG contributed with the conceptualization, writing, reviewing and editing of the current manuscript. The research was conducted by EED supervised by VG and YBA. YBA, EED and VG contributed to funding acquisition. EED developed resources, methodology and the formal analysis.

## Data Access Statement

All data underlying the results are available as part of the article and no additional source data are required.

---

**Figure S1:**
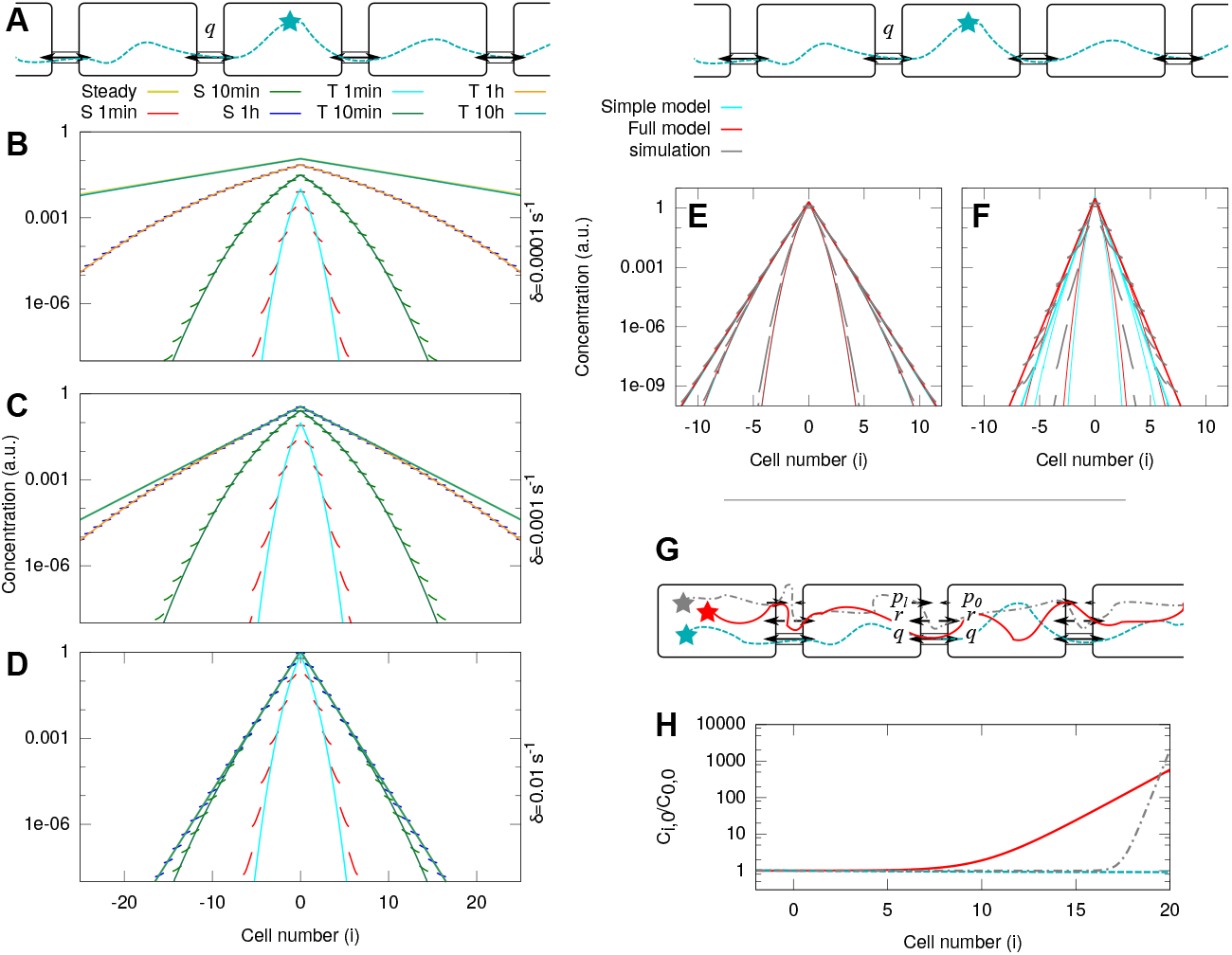
Model verification: analytical and numerical temporal solutions agree for a wide parameter regime. **A**: Setup for **B-F**: Long 1D tissue with production in the middle cell only (with rate *β/*volume = 2*δ* a.u.). Other default parameters: *q* = 1 *µm/s, δ* = 0.001 *s*^−1^, *l* = 100 *µm, D* = 300 *µm*^2^*/s*. **B-D**: Dependence of profile steepness and time scales on decay rate *δ*. In the legend on top of C, simulation profiles are indicated with “S” and analytical predictions with “T”. **E,F**: With high *δ* = 0.1*s*^−1^, analytical predictions with the simplified model (cyan) perform distinctly worse than the full model (red) compared with simulations (gray) as symplasmic permeability *q* decreases from 10 *µm/s* (E) to 1 *µm/s* (F). Only the full model predicts the correct steady state. Profiles at 10 s, 1 min and 30 min. **G**: Setup for **H**: 20 cells 1D tissue with constant production or a fixed concentration at the left of cell 0 (location of source) and a reflecting boundary at the right of cell 20 (organ tip). **H** Comparing steady state gradients for purely apoplasmic transport (gray), purely symplasmic transport (cyan) and mixed transport (red). The addition of symplasmic transport results in a less steep, but much longer gradient and only a minor (∼ 3-fold) decrease of the maximum concentration. Concentrations are plotted relative to cell 0, on a logarithmic scale. (*δ* = 1 ·10^−5^*s*^−1^, *D* = 300 *µm*^2^*/s, q* = 10 *µm/s, p*_0_ = 1 *µm/s, p*_*l*_ = 20 *µm/s, l* = 100 *µm*).

**Figure S2:**
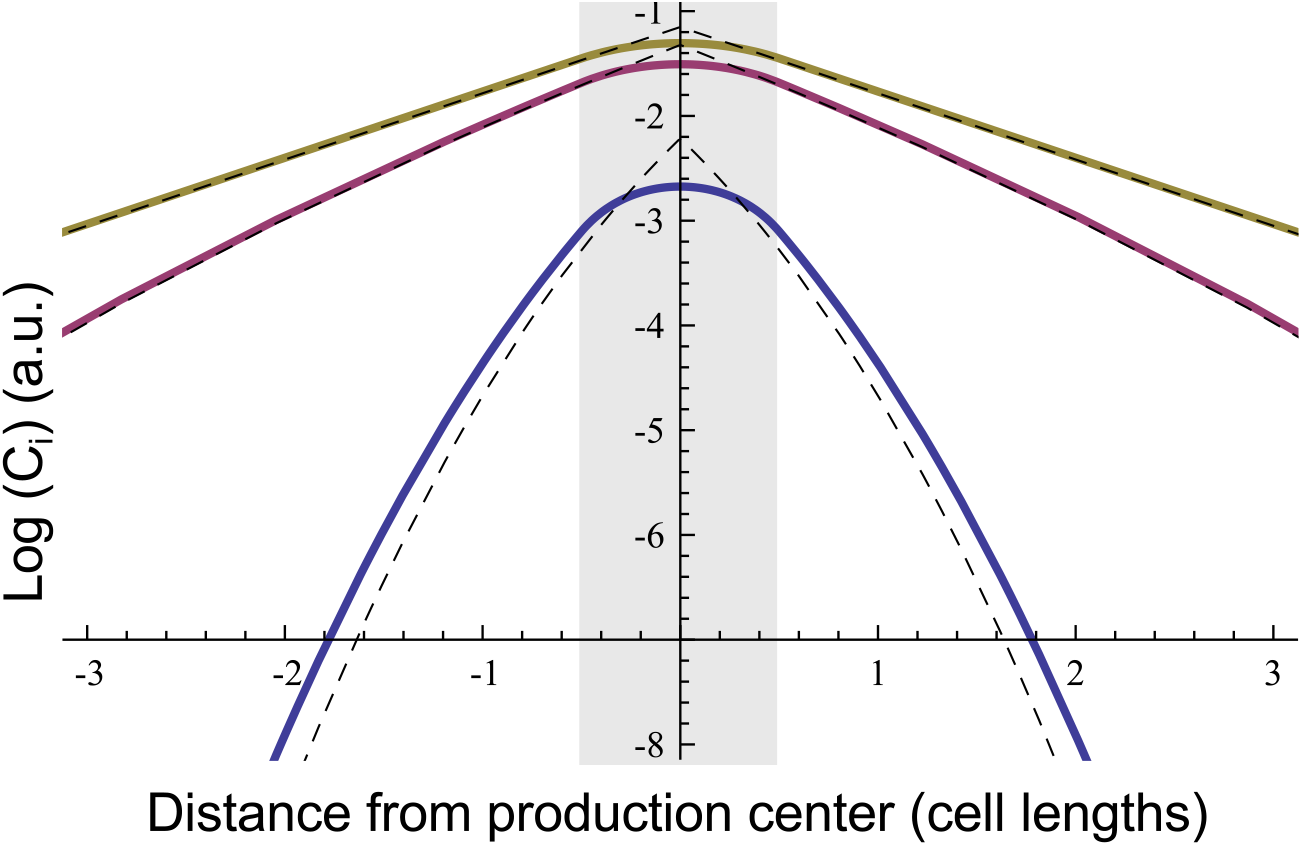
Correction of source shape mostly affects early stages. Theoretical prediction of dynamic behaviour based on full model is normally based on a point source (dashed lines). In the simulation mod el the source is an entire cell. Correcting for this by convoluting the dynamic solution with an entire cell length (shaded grey area) as production source mostly affect the early stages of dispersal (solid blue line) and does not visibly affect the later stages (solid purple and yellow lines). This is a proof of principle with arbitrary parameter values.

**Figure S3:**
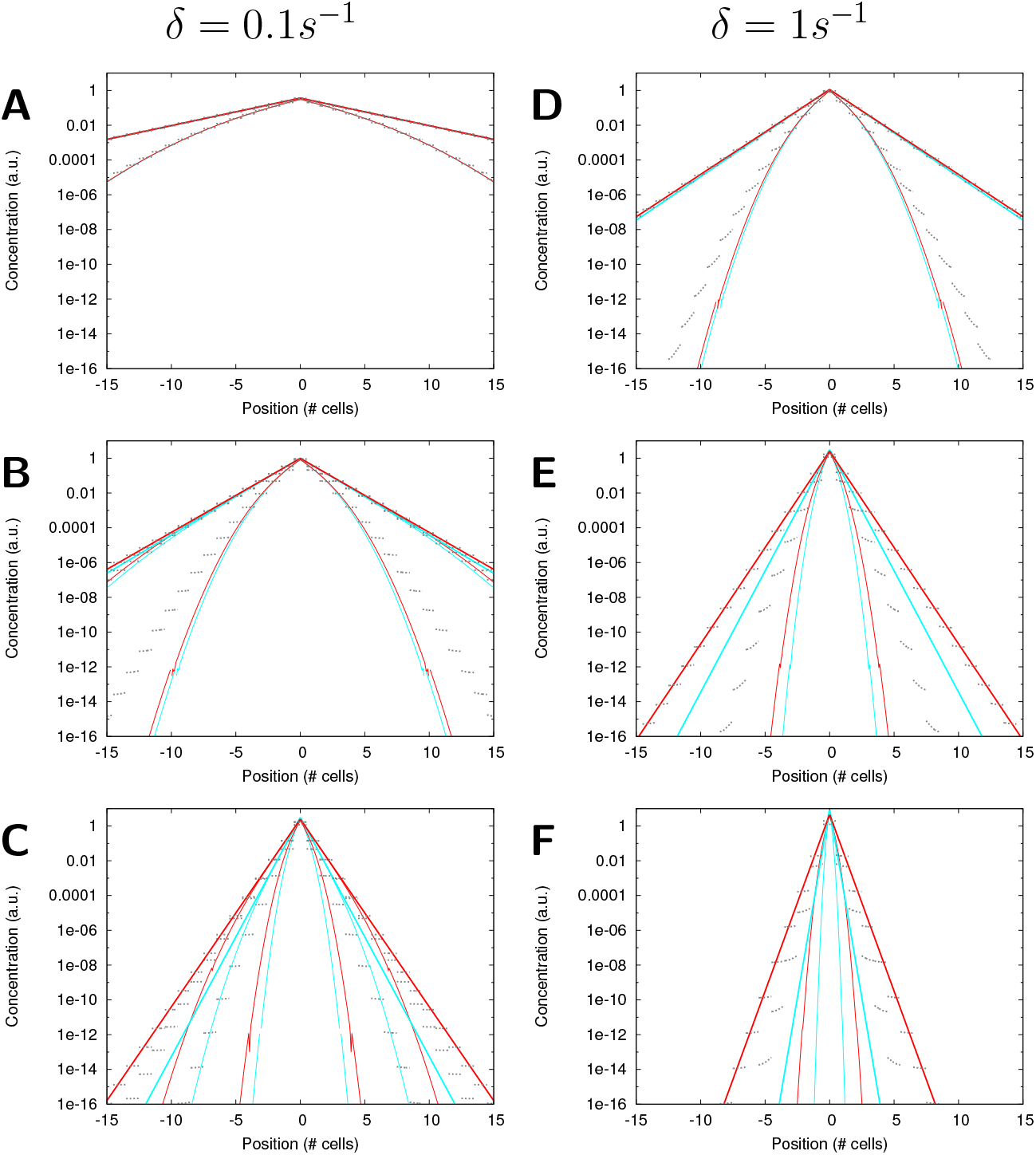
Comparison of full cell based calculations (red/dark) and simplified model (cyan/light) with numerical simulations ((gray) dotted). For a large range of parameters the two yield very similar predictions, e.g. **A** (*q* = 10*µm/s, δ* = 0.1*s*^−1^, *l* = 10*µm, D* = 300*µm*^2^*/s*). The predictions start to deviate towards high (or very high) *δ*, particularly when combined with low *q* (towards the bottom of the figure) and/or long cells (Figure S4). Under such circumstances, the full model consistently performs better than the simple model and only the full model correctly predicts the steady state distribution. A,B: *q* = 10 *µm/s*. C,D: *q* = 1 *µm/s*. E,F: *q* = 0.1 *µm/s*. A,C,E: *δ* = 0.1*/s*, B,D,F: *δ* = 1*/s*. All panels: *l* = 10 *µm, D* = 300*µ m*^2^*/s*.

**Figure S4:**
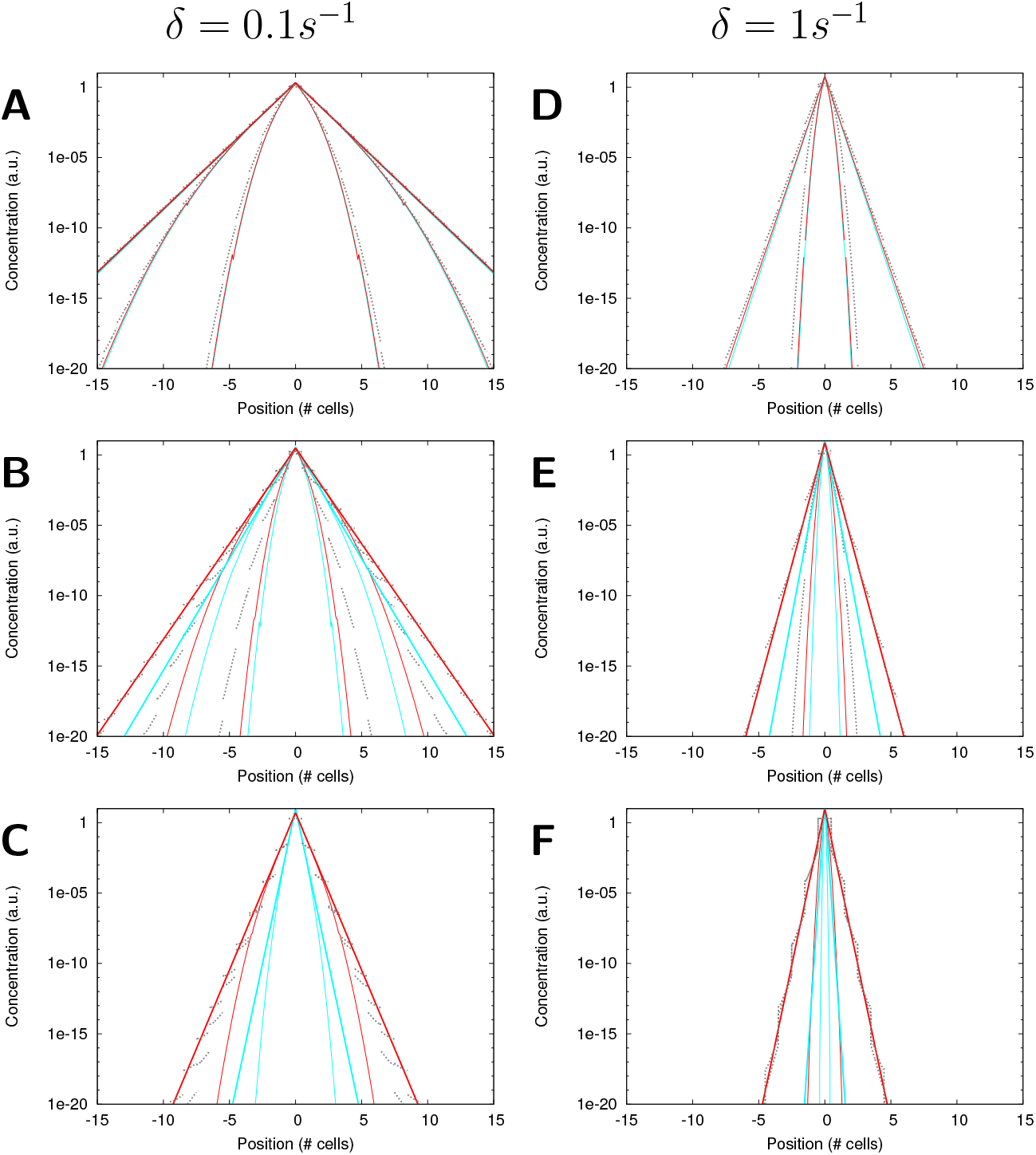
Comparison of full cell based calculations (red/dark) and simplified model (cyan/light) with numerical simulations ((gray) dotted). As Figure S3 but with long cells (*l* = 100 *µm*). For a large range of parameters the two yield very similar predictions, e.g. **A** (*q* = 10*µm/s, δ* = 0.1*s*^−1^, *l* = 100*µm, D* = 300*µm*^2^*/s*). The predictions start to deviate towards high (or very high) *δ*, particularly when combined with low *q* (towards the bottom of the figure) and/or long cells (short cells in Figure S3). Under such circumstances, the full model consistently performs better than the simple model and only the full model correctly predicts the steady state distribution. A,B: *q* = 10 *µm/s*. C,D: *q* = 1 *µm/s*. E,F: *q* = 0.1 *µm/s*. A,C,E: *δ* = 0.1*/s*, B,D,F: *δ* = 1*/s*. All panels: *l* = 100 *µm, D* = 300*µ m*^2^*/s*.

**Figure S5:**
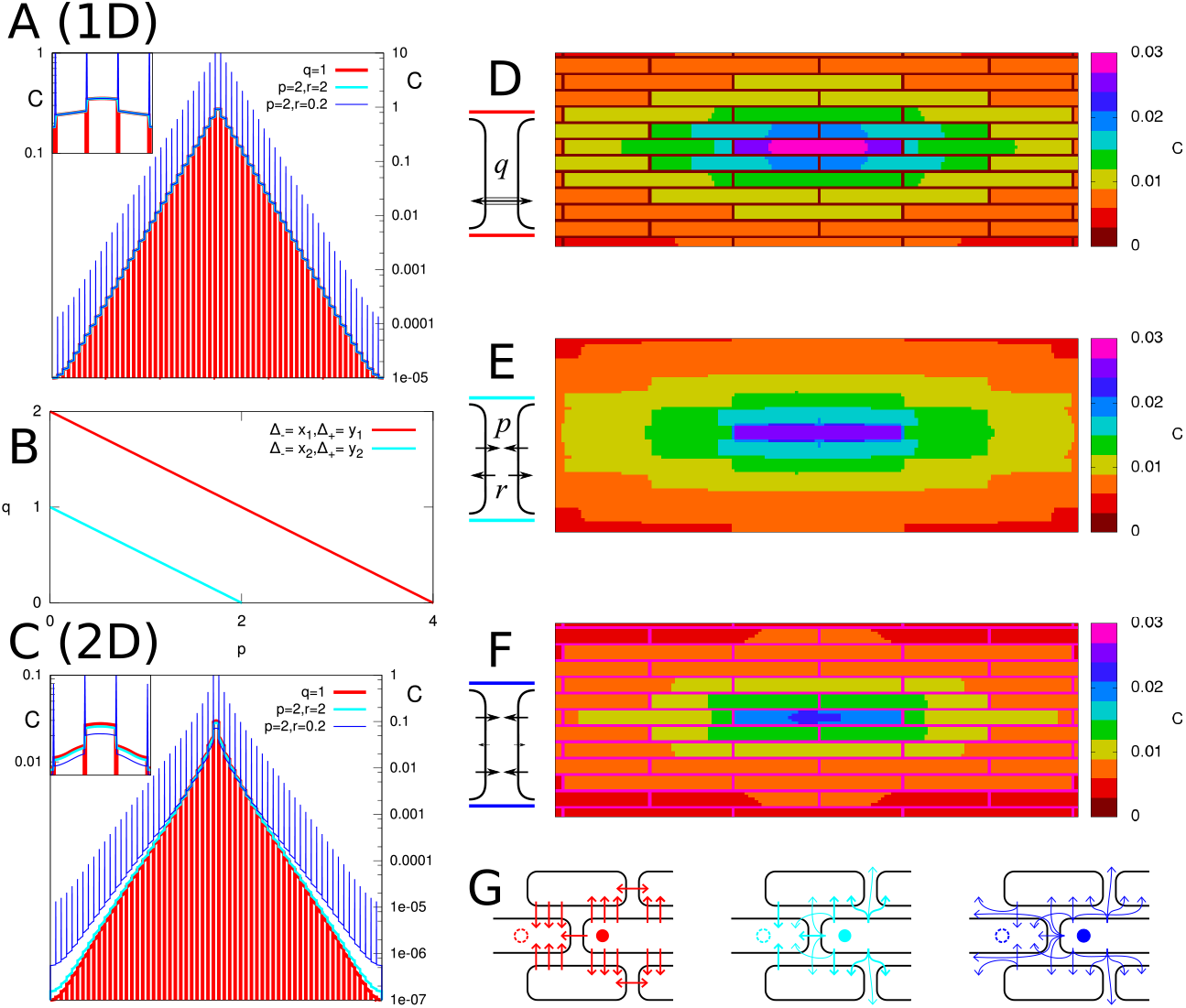
Identical steady state profiles possible in 1D, but not in 2D. Signal production in the central cell. **A**: 1D steady state profiles: for parameters satisfying *y* = 2*p* + *q* (see **B**), only the wall concentrations differ between symplasmic (red) and symmetric apoplasmic (cyan: equal effective influx and efflux permeabilities *r* = *p*, blue: reduced effective influx permeability *r* = *p/*10) transport. **C**: with the same parameters on a 2D tissue (brick alignment: see **D-F**) the intracellular concentrations are no longer the same (though profiles remain similar), because it is possible to bypass cells through the apolast, as sketched in **G**. Parameters: *D* = 300 *µm*^2^*s*^−1^, *l* = 100 *µm, δ* = 0.001*s*^−1^, *β* = 0.002[*C*]*s*^−1^, *q* = 1 *µm/s* (**D**: symplasmic only), *p* = 2 *µm/s, r* = 2, 0.2 *µm/s* (**E,F**, respectively: both apoplasmic only).

**Figure S6:**
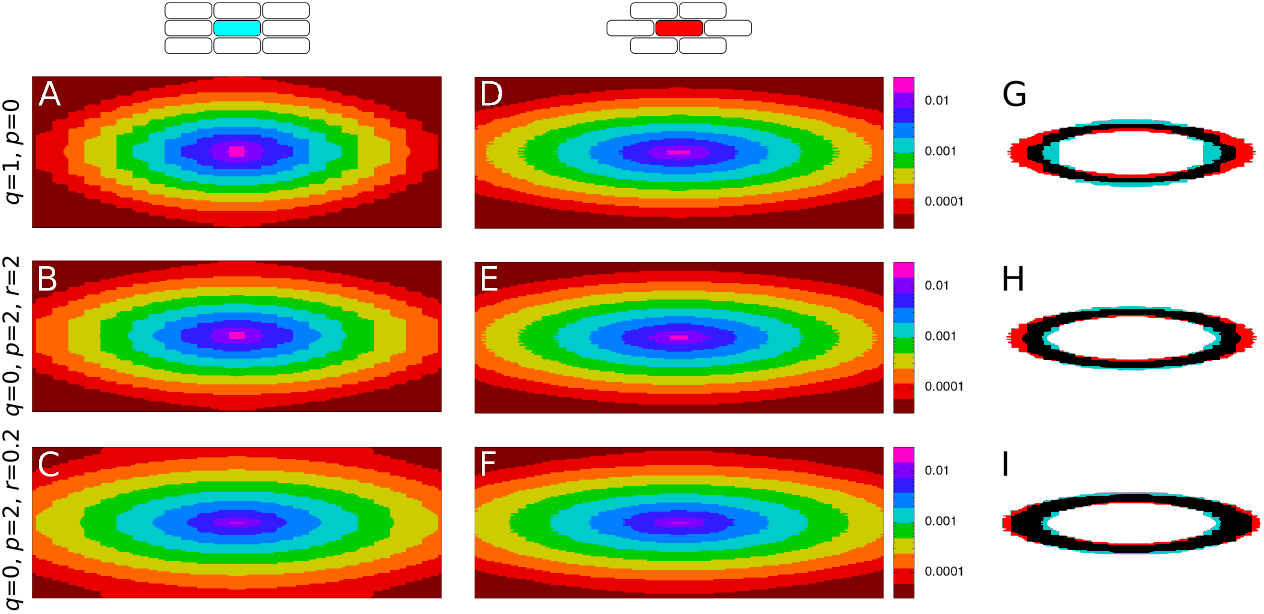
Symplasmic transport is more sensitive to tissue layout than apoplasmic transport. **A-F**: Concentration heat maps excluding cell wall concentrations. Production takes place in the central cell. **A-C**: Square alignment. **D-F**: Brick alignment. **G-I**: Concentrations are indicated with different colors using a step function. An overlay of a ring representing the same concentration range for both alignments is shown on the right (cyan: “square”, left; red: “brick”, middle; black: overlap). Differences are largest for symplasmic transport (top row), indicating that this transport mode is most sensitive to tissue layout. **A,D,G**: Only symplasmic transport (*q* = 1*µm/s*). **B,E,H**: Only apoplasmic transport, with high uptake rate (*p* = 2*µm/s, r* = 2*µm/s*). **C,F,I**: Only apoplasmic transport, with low uptake rate (*p* = 2*µm/s, r* = 0.2*µm/s*). Parameters: *D* = 300 *µm*^2^*s*^−1^, *l* = 100 *µm, δ* = 0.001*s*^−1^, *β* = 0.002[*C*]*s*^−1^.

**Figure S7:**
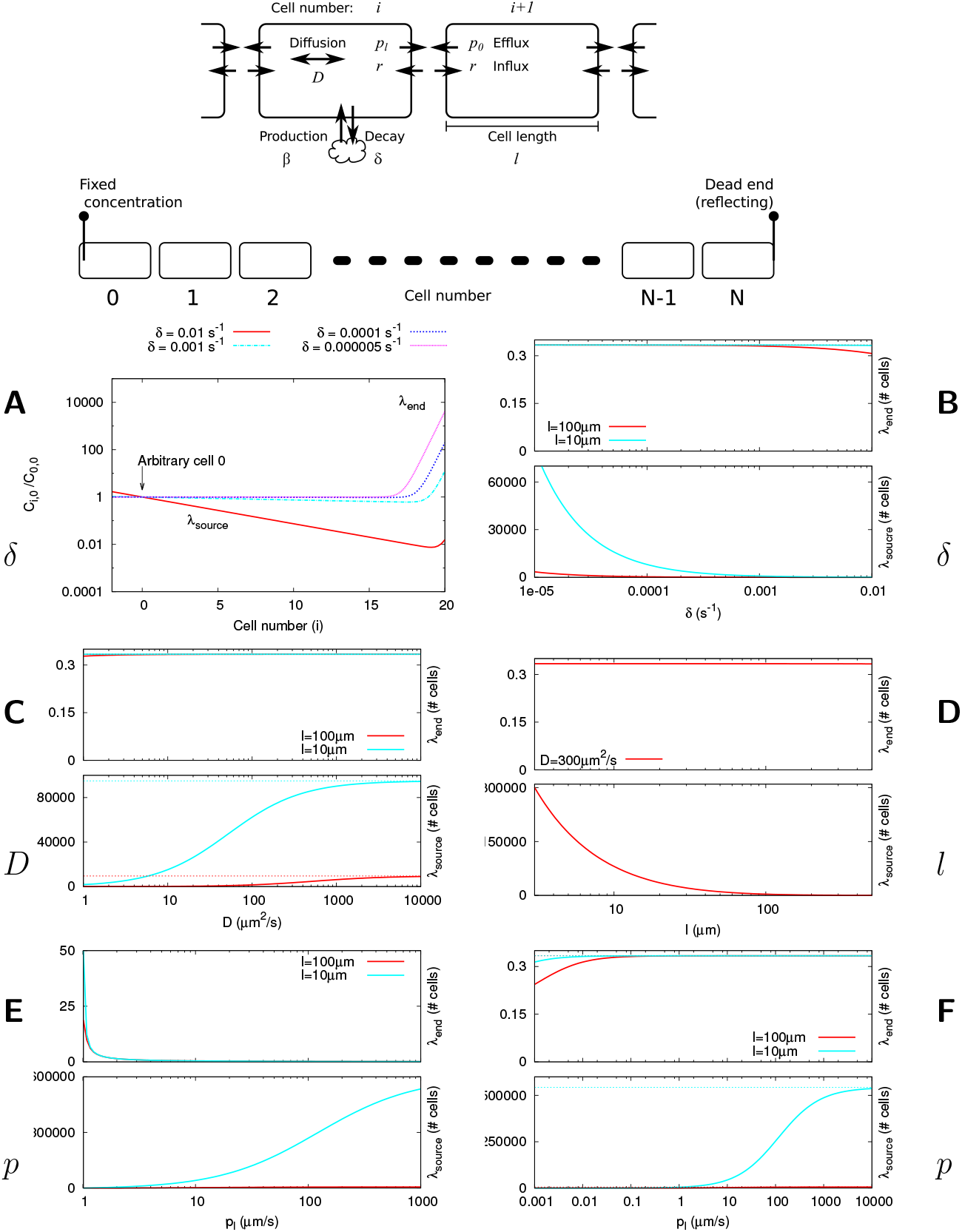
Purely apoplasmic transport (*q* = 0). **A**: example profiles for different values of *δ*. Note that with high *δ* only little substance reaches the far end. We introduce two characteristic lengths (*λ*_*source*_ and *λ*_*end*_, both expressed in number of cells). Note that a “heaping up” effect at the far end is only possible if *λ*_*source*_ is large. **B-E**: dependence of *λ*_*source*_ and *λ*_*end*_ on single model parameters. **F**: dependence of *λ*_*source*_ and *λ*_*end*_ on *p*_*l*_, with a fixed ratio *p*_*l*_*/p*_0_ = 20. Default values: *p*_*l*_ = 20*µms*^−1^, *p*_0_ = 1*µms*^−1^, *D* = 300*µm*^2^*s*^−1^, *l* = 100*µm, δ* = 0.00001*s*^−1^.

**Figure S8:**
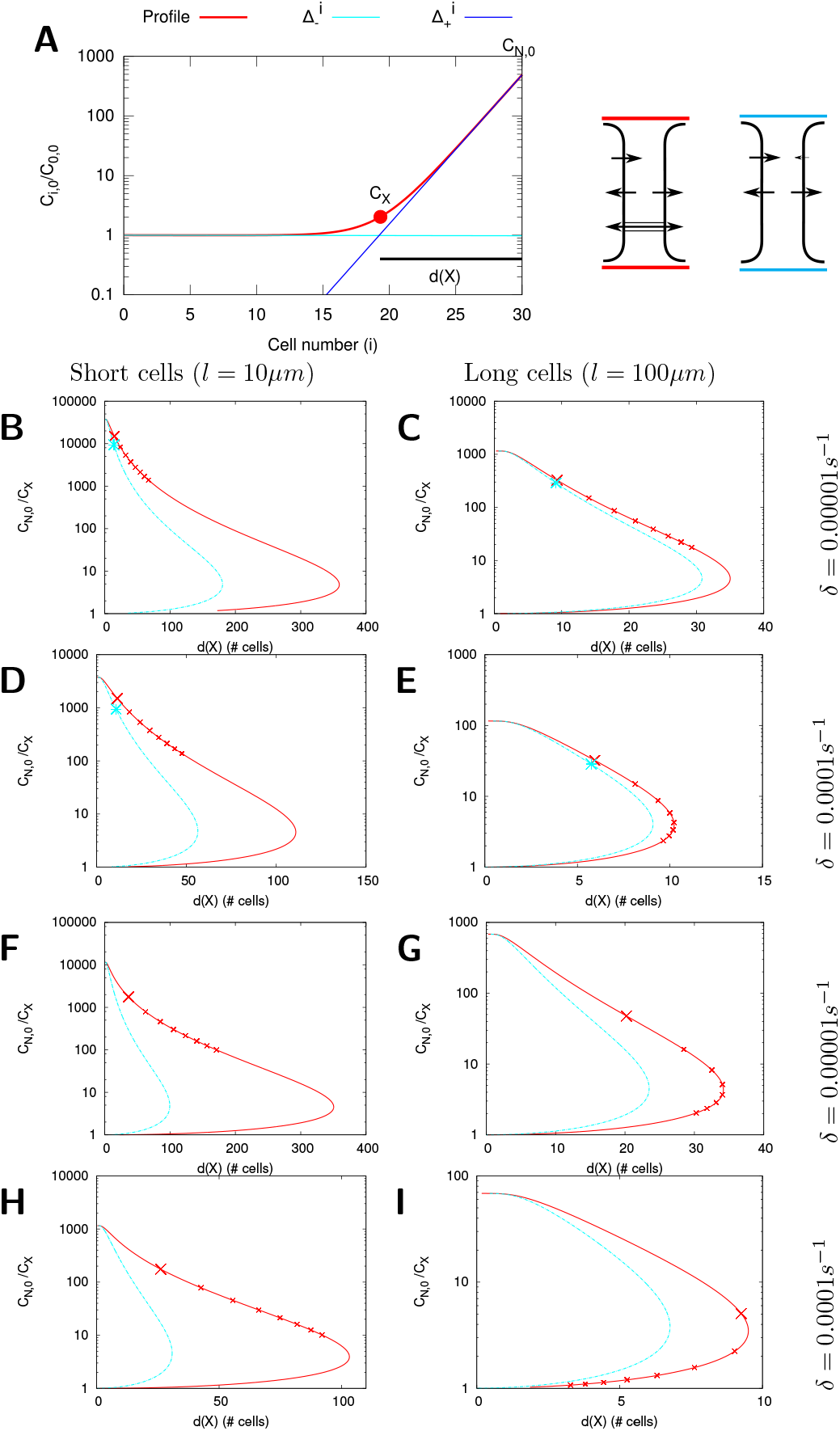
Symplasmic vs. apoplasmic backflux for different parameters. Curves show the length of the informative gradient (*d*(*X*)) and relative end-concentration (*C*_*N*,0_*/C*_*X*_), as explained in **A**, for symplasmic (red curves) and apoplasmic (cyan curves) backflux. The crosses on the symplasmic curves occur every 10*µm/s* until 80*µm/s*, showing how progression along this curve slows down with increasing *q*. Parameters: *p*_*l*_ = 20*µm/s* (**B-E**) or *p*_*l*_ = 5*µm/s* (**F-I**), *D* = 300*µm*^2^*/s, r* = 20*µm/s*.

**Figure S9:**
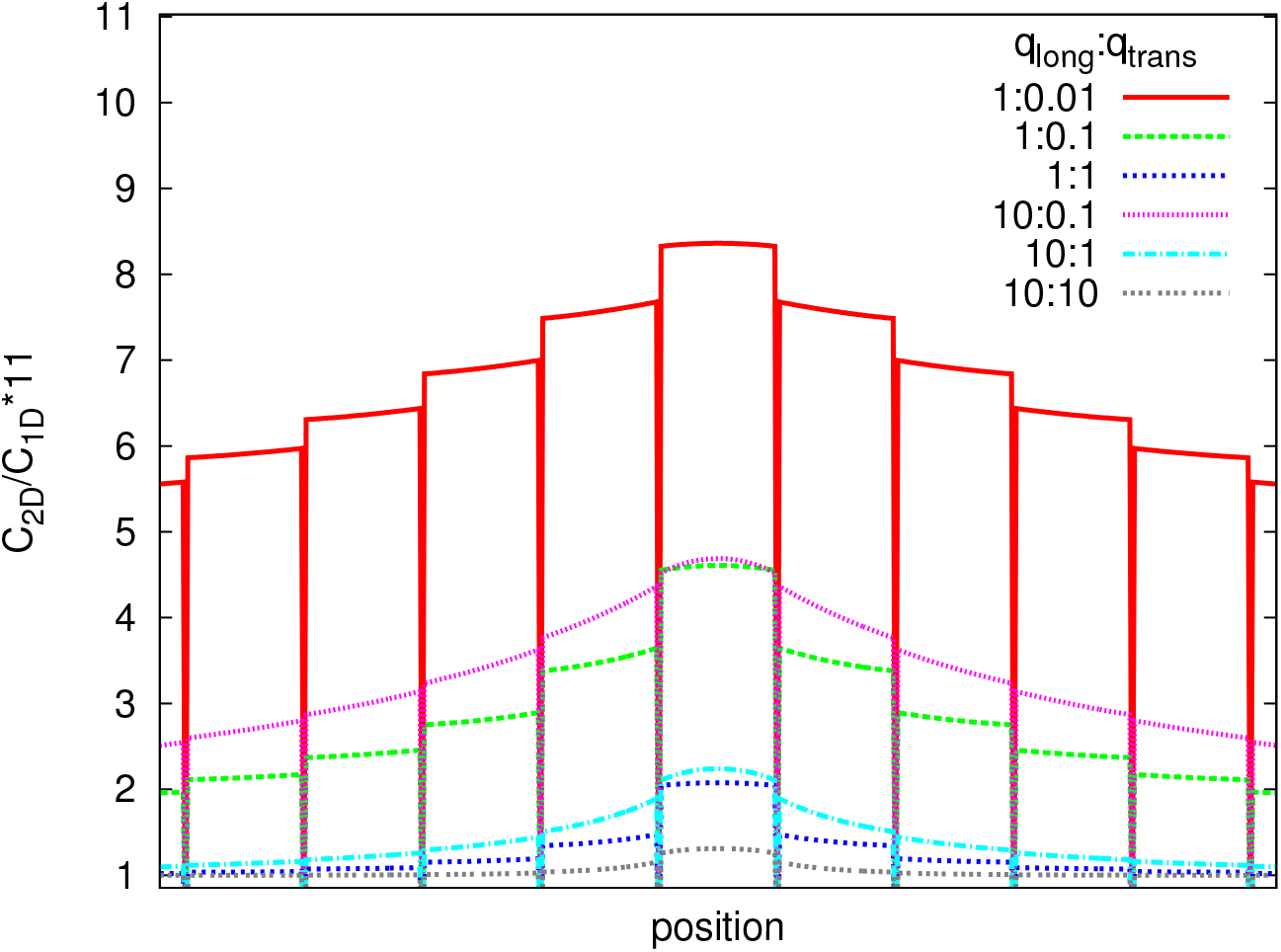
Comparison of impact of *q*_*l*_ and *q*_*t*_ on how, moving away from the source, the profile relaxes to a 1D profile with the same parameters (*q*_1*D*_ = *q*_*l*_). Curves are calculated based on Figure 4A-F: concentration in 2D in the cell file containing the producing cell divided by the concentration in 1D, multiplied by 11. As a result of this multiplication, a value of 1 signifies local equivalence to the corresponding 1D profile. Note that the spatial relaxation to the 1D equivalent for equal *q*_*t*_ is more similar than for equal *q*_*l*_ : *q*_*t*_.

**Figure S10:**
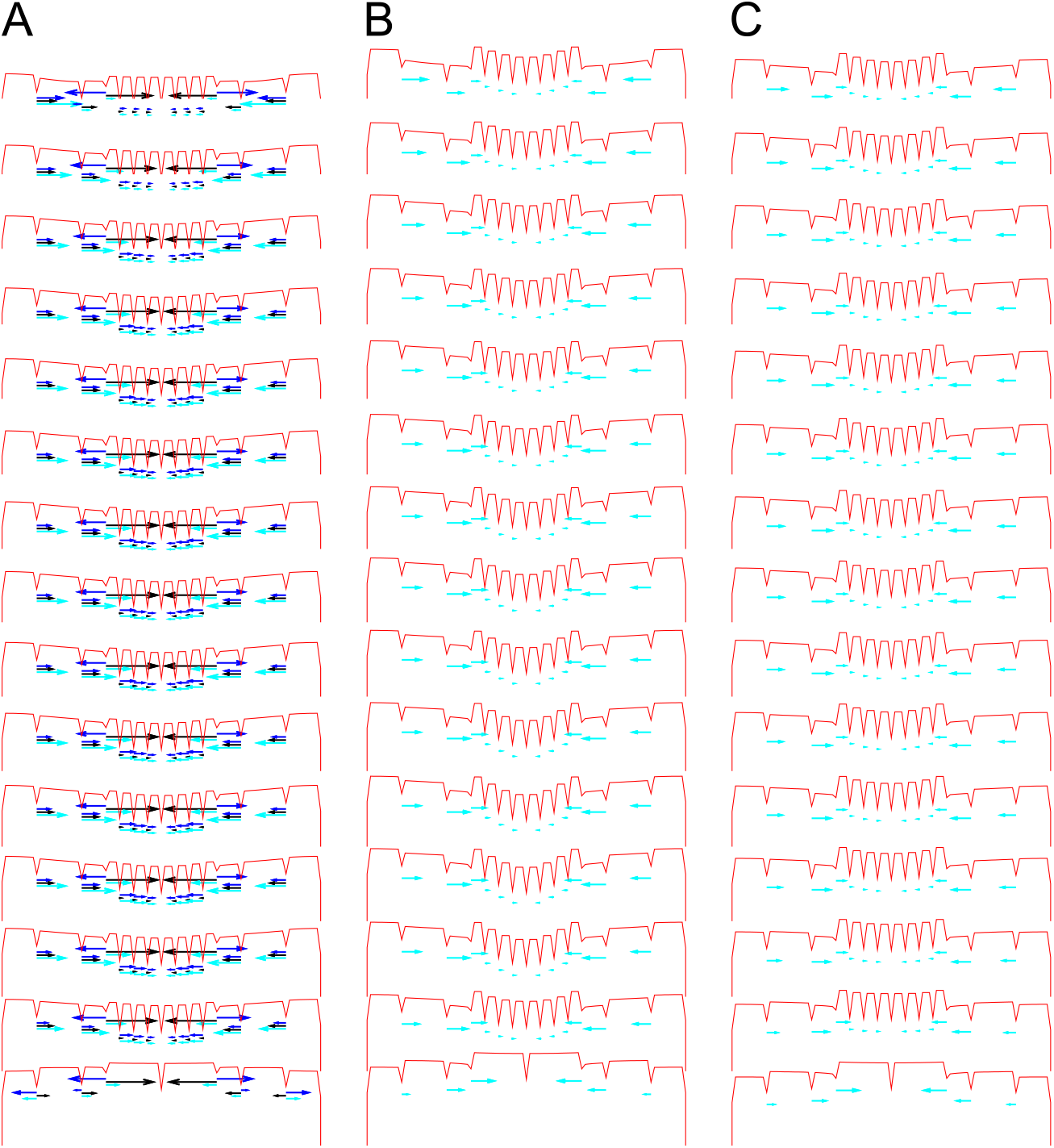
Impact of symplasmic transport on root gradients. Radial concentration profiles (red) for all MZ cell layers in **Figure 5C-E**. Net radial fluxes are indicated by cyan arrows, with arrow length proportional to the relative flux. Where symplasmic transport is possible in the radial direction (**A**), blue arrows show the symplasmic and black arrows the apoplasmic components of the fluxes. Note that the net inward fluxes are larger in **A** than in **B** and **C**. Roots based on Laskowski et al. [42], with slightly altered cell sizes. Symplasmic transport is possible in the radial direction (**A**) or longitudinal direction (**C**) of the root, or not at all (**B**). No symplasmic transport in the columella (*C* = 0). *δ* = 5 · 10^−6^*s*^−1^.

**Figure S11:**
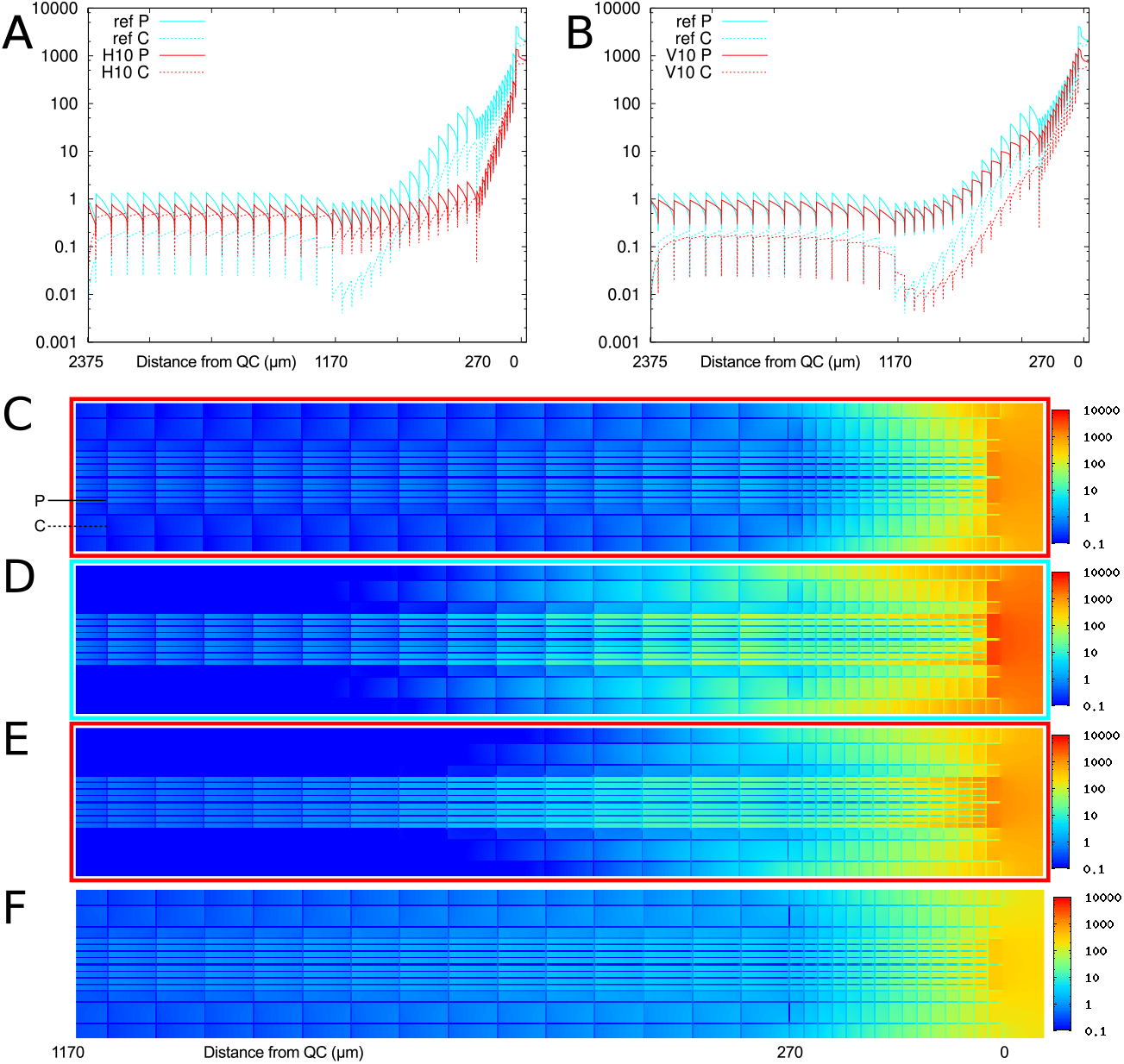
Impact of symplasmic transport on root gradients. symplasmic transport is possible in the radial direction (**A,C**) or longitudinal direction (**B,E**) of the root (red profiles). Control roots without symplasmic transport (**D**) in cyan. **F**: Profile with both radial and longitudinal transport looks more similar to radial than longitudinal only. Numbers below graphs and pictures indicate distance from the QC. Symplasmic transport also occurs over all columella walls (*C* = 10 *µm/s*).

## A Mathematical derivations

### A.1 1D analytical model: coordinate system

We track both concentration (*C*_*i,x*_) and flux, in the positive direction, (*J*_*i,x*_) with subcel-lular precision. The cell number (*i*) and position (*x* ∈ [0, *l*]) are indicated with subscripts.

### A.2 1D steady state profile for purely diffusive symplasmic transport

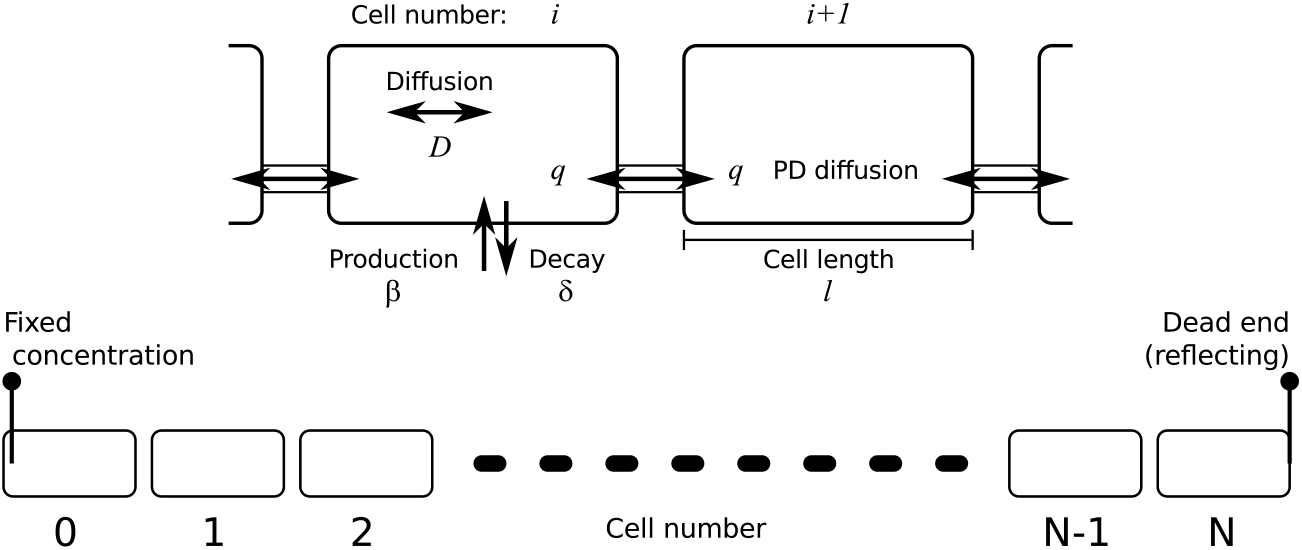

With pure symplasmic transport, the flux over a wall is given by:

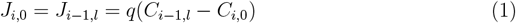

To link one end of the cell to the next, we compute the intracellular concentration profile. With (only) decay with rate *δ* this profile *f* (*x, t*) must obey:

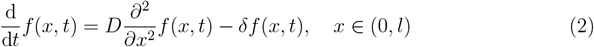

Which has the general steady state solution

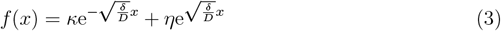

with *κ* and *η* constants. For a given cell *i*:

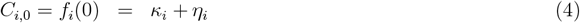

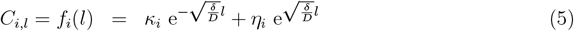

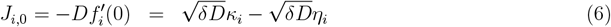

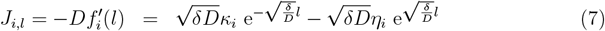

Choosing the least complicated equations we start from *C*_*i*,0_ and *J*_*i*,0_:

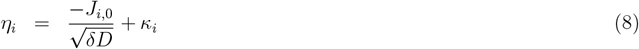

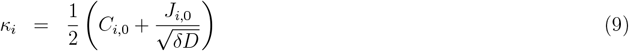

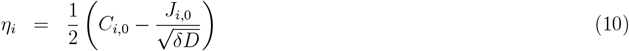

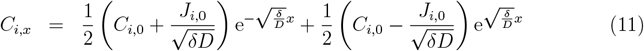

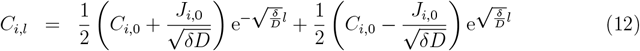

Matter is conserved, so the flux over a given wall is the same as the flux over the previous wall minus the decay in that cell:

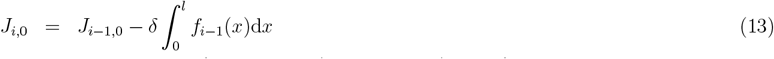

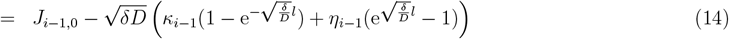

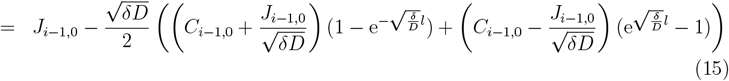

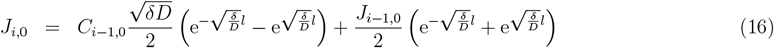

We introduce the ratios

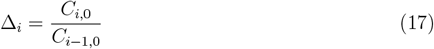

With these we can search for basis solutions, i.e. Δ_*i*_ = Δ_*i*+1_ = Δ. Introducing the short hand notations

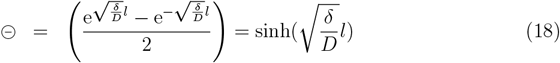

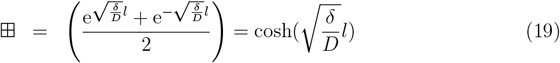

we can rewrite equations 16 and 1:

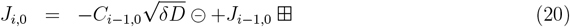

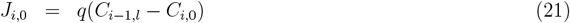

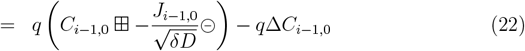

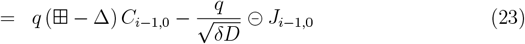

So

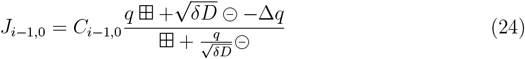

At the same time

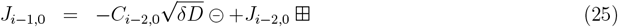

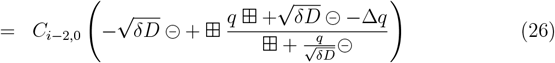

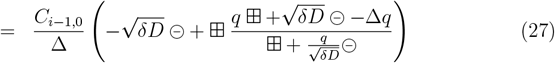

Combining Equation 24 and 27, and solving for Δ we obtain two basis solutions:

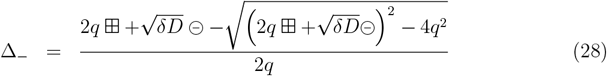

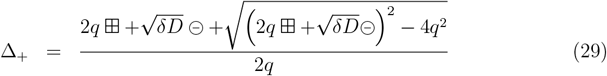

This means that the general steady state profile must be a linear combination of these two solutions:

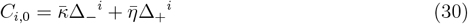

We use a no flux boundary at the last cell (*N*), so at steady state:

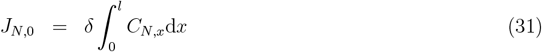

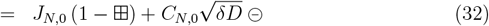

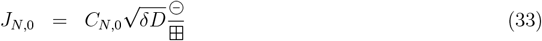

and

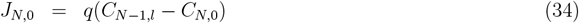

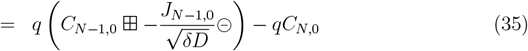

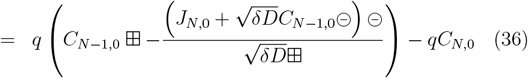

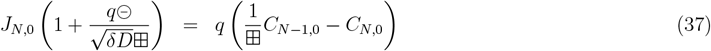

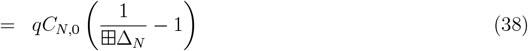

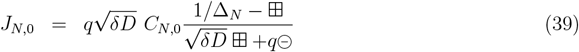

Combining Equation 33 and 39, and solving for Δ_*N*_ :

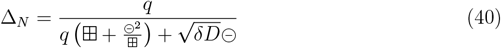

We now have

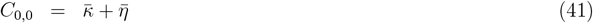

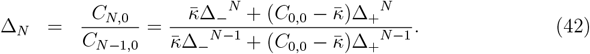

Leaving out the middle expression and solving for 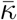 and 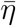, we get

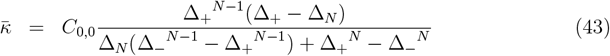

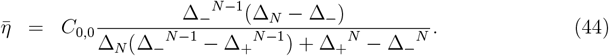

#### A.2.1 Characteristic length

It is straightforward to show that Δ_−_ *<* 1 and Δ_+_ *>* 1. This means that in the limit of *N* → ∞, 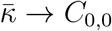 and 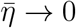, so

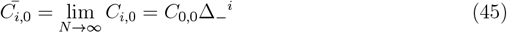

This we can rewrite as a single negative exponential:

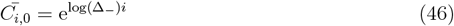

In the literature of animal morphogen gradients the characteristic length of a gradient is defined as *λ* in

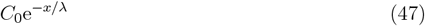

with *C*_0_ the concentration at the source and *x* a spatial coordinate. On an infinite line of cells our profile thus has the characteristic length

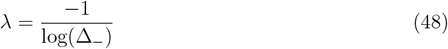

in number of cells or

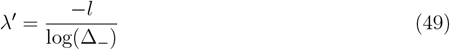

in real length (*µm*).

#### A.2.2 Useful limits

To verify the formulas, we compute a few physically meaningful limits. By taking the limit for *D* → ∞ (using l’Hôpital’s rule) we arrive at the expressions that can be found without taking into account the intracellular gradients.

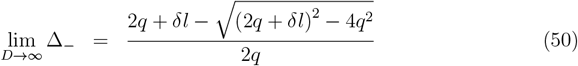

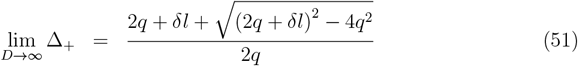

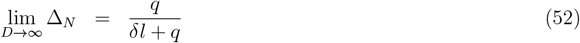

If we consider the walls as interfaces of vanishing thickness and let the diffusive permeability *q* → ∞ we obtain:

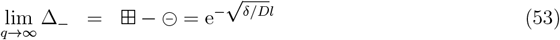

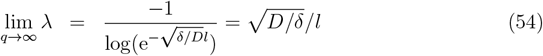

for position expressed in cell number. This can be changed to actual length by multiplying by the cell length, obtaining 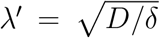, exactly the characteristic length found for diffusion/decay gradients.

Inspired by these limits, we could write an “effective diffusion coefficient” *D*_*eff*_ from the characteristic length:

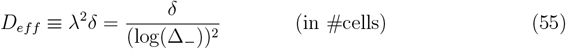

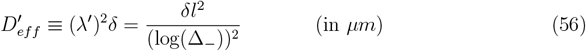

These are dependent on *δ*, which seems odd. Taking the limits for *δ* ↓ 0, we obtain:

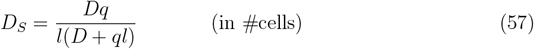

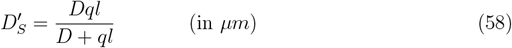

The latter expression is also found by [15], describing effective diffusion in an animal tissue (1D). We therefore tested if these formulas were also useful for *δ >* 0 to map our system (in a coarse-grained way) to ordinary diffusion/decay system. This worked surprisingly well (e.g. Figure S2), but the more elaborate formulas performed better for high *δ* in combination with low *q* and long cells (high *l*) (e.g. Figure S4).

### A.3 1D: time resolved solution for purely diffusive symplasmic transport (approximation)

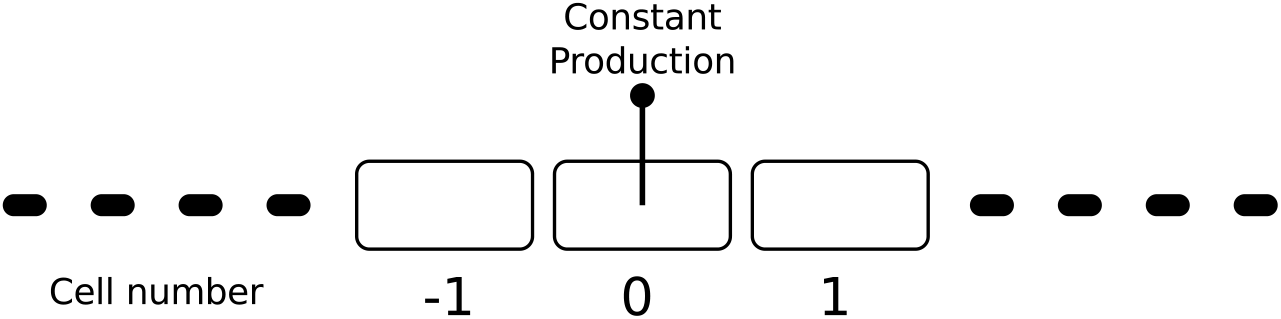

Assume an infinite file of cells, *k* ∈ (−∞, ∞), with production only in the middle cell (*k* = 0), with rate *β*. In an attempt to obtain the full time dependent solution, we exploit that the steady state can be rescaled to a homogeneous diffusion problem. We assume that all matter produced in cell 0 at time *t* will spread outward following a Gaussian profile with *σ*^2^ = 2*αt, α* some yet unknown constant. Including homogeneous decay with a constant rate *δ*, this will be:

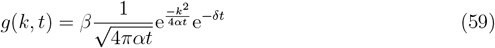

Integrating over all time till moment *T* :

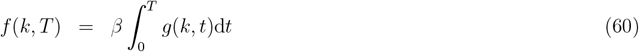

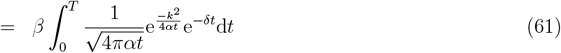

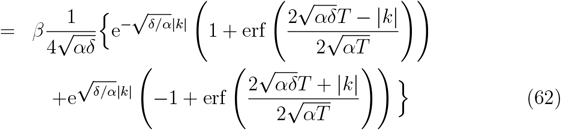

As lim_*x*→∞_ erf(*x*) = 1,

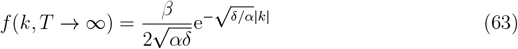

We also know that without boundaries, similar to Equation 46, the steady state profile must follow

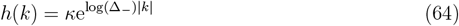

with *κ* some constant and

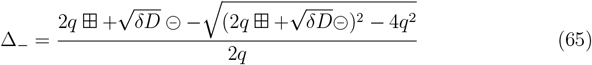

We can thus solve for *α*:

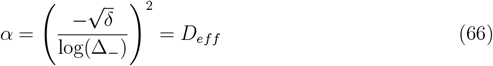

So

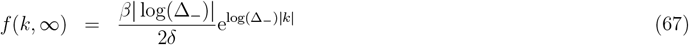

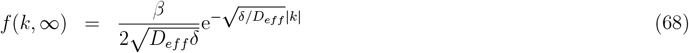

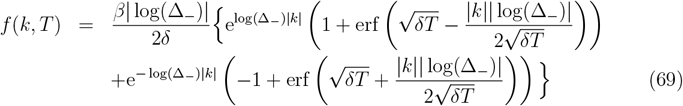

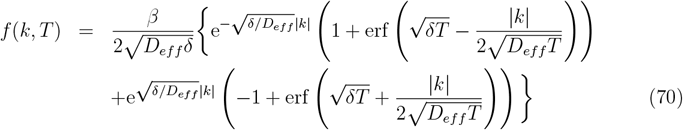

Note that in this approach we implicitly assume that within each cell, the concentration profile follows a steady state distribution (quasi steady state assumption). This is, of course, only true in the actual steady state. If, however, the deviations from the steady state profile are sufficiently small, Equation 69 will be a good approximation for the tissue profile. Moreover, the difference between this profile and the actual non-steady state profile (obtained from numerical simulations with subcellular precision) will decrease with time and will eventually vanish (Figure S3, S4). A partial correction for the tails can be obtained by taking into account the actual size (1 cell) of the source. This results in fatter tails in early stages and converges to the same steady state profile (except for the source cell, which actually is an improvement too) for the tissue (Figure S2). Unfortunately, there is no analytical expression for the corrected solution, so one has to resort to numerical integration.

### A.4 1D steady state profile for combined symplasmic transport and apoplasmic transport

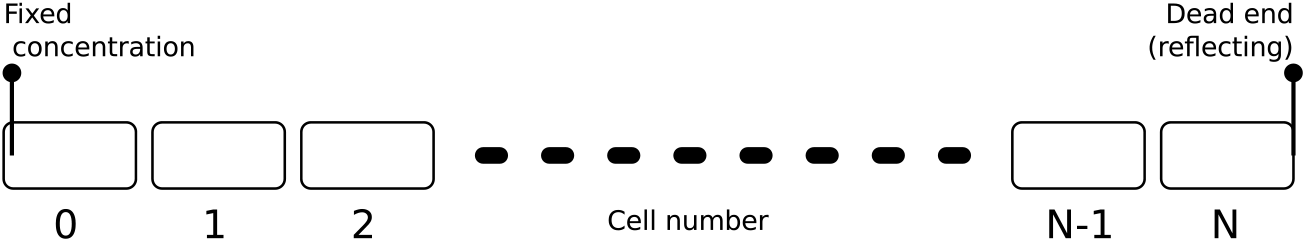

*This is a single derivation for the two processes, symplasmic and apoplasmic transport, combined. Equations for either process can be obtained by setting the parameters for the other process to zero. For apoplasmic transport this is only possible with efflux on both sides (possibly very small on one side), otherwise the resulting formulas contain a division by zero. In essence the derivation is the same as for only symplasmic transport (appendix A.2), except for a more complicated expression for the flux over the wall, naturally affecting the resulting expressions*.

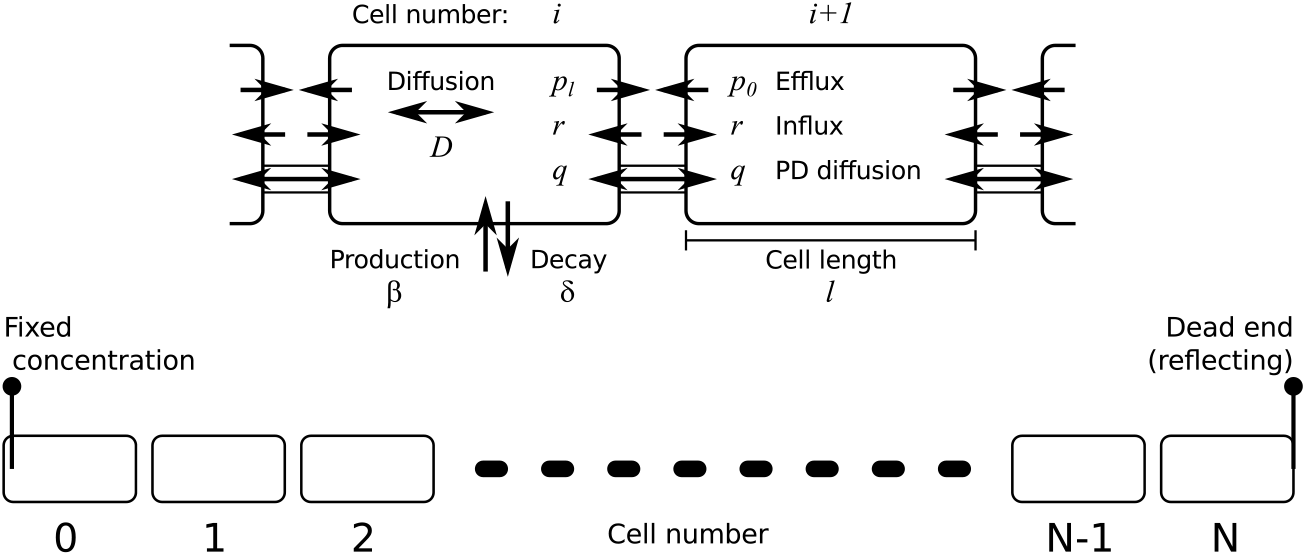

In the setting we chose, only diffusion and decay take place inside a cell. That means that the intracellular profiles are the same as before:

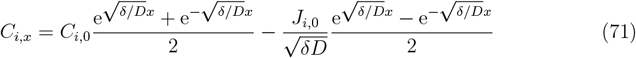

So, using the shorthand notation introduced before (equations 28 and 29):

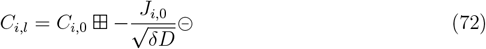

The fluxes over the wall follow

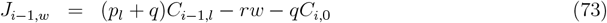

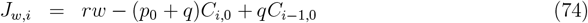

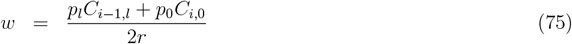

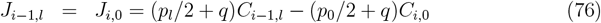

Moreover,

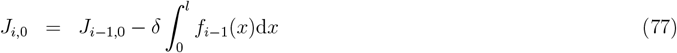

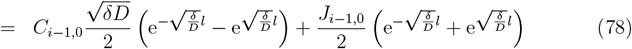

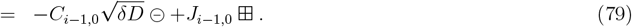

We again compute the ratios

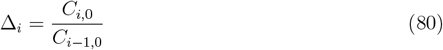

and search for basis solutions obeying Δ_*i*_ = Δ_*i*+1_ = Δ ∀*i*.

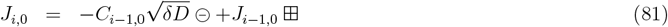

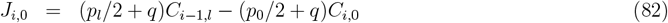

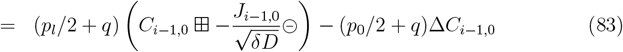

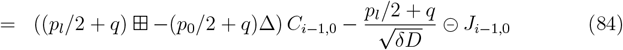

So, combining Equation 79 and 84,

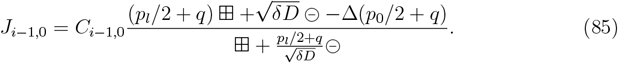

Similarly, for the preceding cell holds:

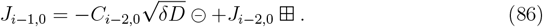

Substituting Equation 85:

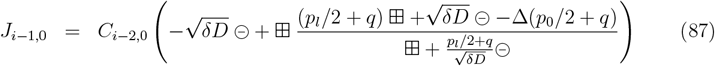

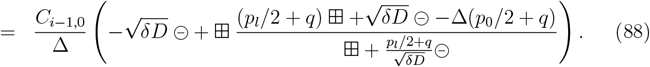

Combining Equation 85 and 88, and solving for Δ:

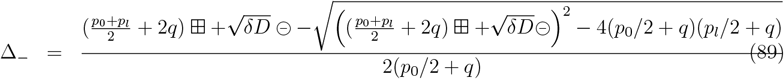

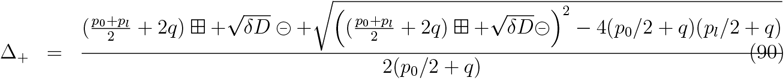

At steady state we have because of the no flux boundary of the last cell:

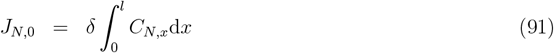

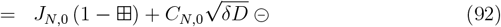

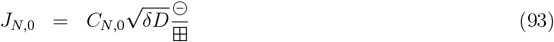

and

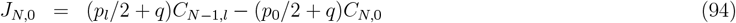

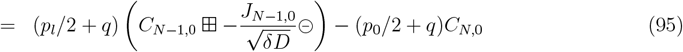

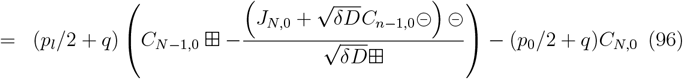

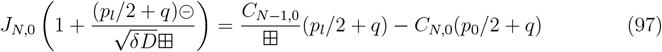

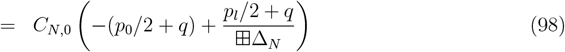

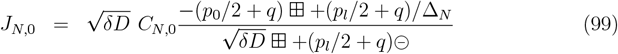

Combining Equation 93 and 99 and solving for Δ_*N*_ :

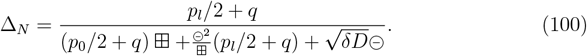

We now have:

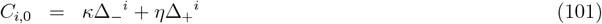

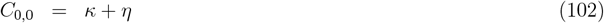

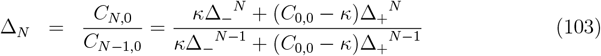

So,

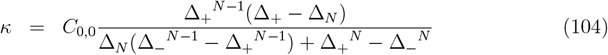

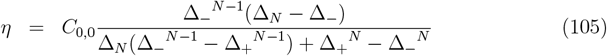

Considering that Δ_−_ *<* 1 *<* Δ_+_, it is not hard to see that for *N* sufficiently large, *κ* ≈ *C*_0,0_ (slightly smaller than *C*_0,0_) and that *η* ≈ 0 (but strictly positive). In other words, in a sufficiently long “tissue” the near end (*i* small) of the profile will be dominated by *κ*Δ_−_^*i*^. With directed apoplasmic transport (*p*_*l*_ *> p*_0_), the far end (*i* ≈ *N*) will be dominated by *η*Δ_+_^*i*^.

Note that (using l’Hôpital’s rule)

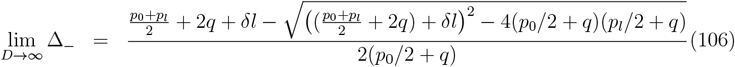

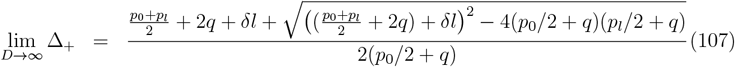

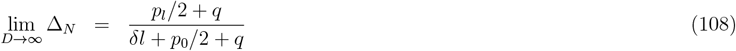

i.e. the same as can be calculated without intracellular gradients.

#### A.4.1 Length of the “informative gradient”

We define the distance *d*(*X*) as the distance between the far end and the point where both parts contribute equally to the solution (i.e. *X* in *κ*Δ_−_^*X*^ = *η*Δ_+_^*X*^):

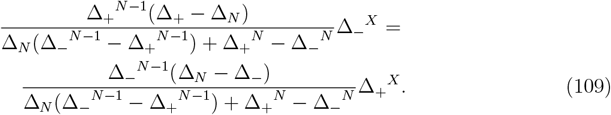

Solving for *X* yields:

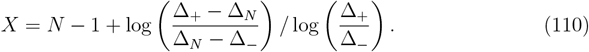

So,

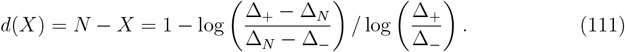

Note that this distance *d*(*X*) is independent of the number of cells. This means that the length of the informative gradient is independent of the total tissue length (provided that it is long enough, i.e. *N > d*(*X*)).

### A.5 1D: Reconstructing intracellular gradients and local fluxes from analytical steady state profiles

The tissue scale profiles calculated in appendices A.2 and A.4 yield the concentrations *C*_*i*,0_ at the “upstream” side of each cell. This contains enough information for reconstructing all intracellular profiles and intercellular fluxes.

#### A.5.1 Intracellular profiles

Inside the cell we have:

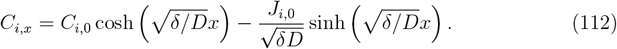

Combining this with Equation 85, found during the derivation in A.4, with Δ_*i*+1_ substituted by its definition

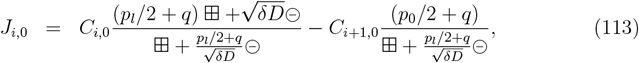

we get:

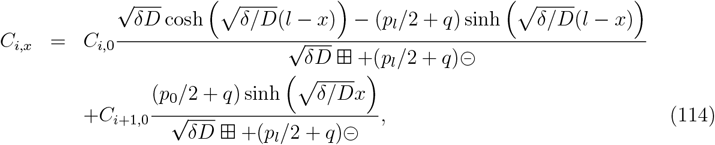

which describes the profile within a cell, based on the concentration at its 0-end and the concentration at the beginning (0-end) of the next cell.

#### A.5.2 Intercellular fluxes

Using the above formula, the concentration at the far end of cell *i* is:

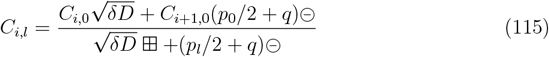

The total flux over the walls at both ends are given by:

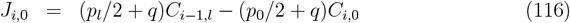

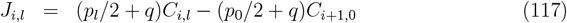

Filling in the expression of *C*_*i,l*_ in *J*_*i,l*_ to obtain the full expression:

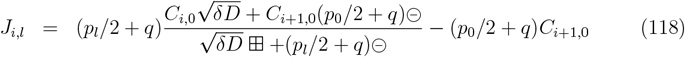

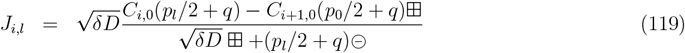

Of this, the net flux through the plasmodesmata is:

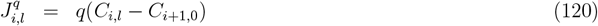

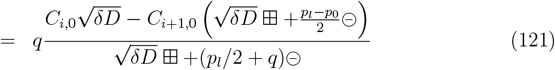

Note that this is not the same as simply dropping *p*_*l*_ and *p*_0_ from the numerator of the full flux equation *J*_*i,l*_.

Similarly the forward apoplasmic flux is:

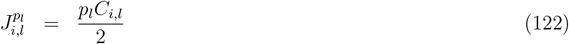

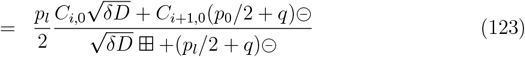

and the reverse apoplasmic flux:

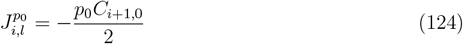

For completeness we write down the same fluxes for the near end of the cell:

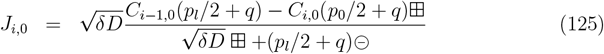

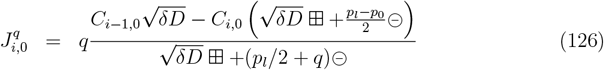

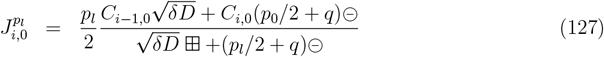

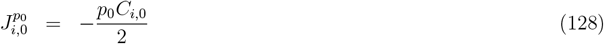

## B Overview of mathematical symbols

**Table S1:**
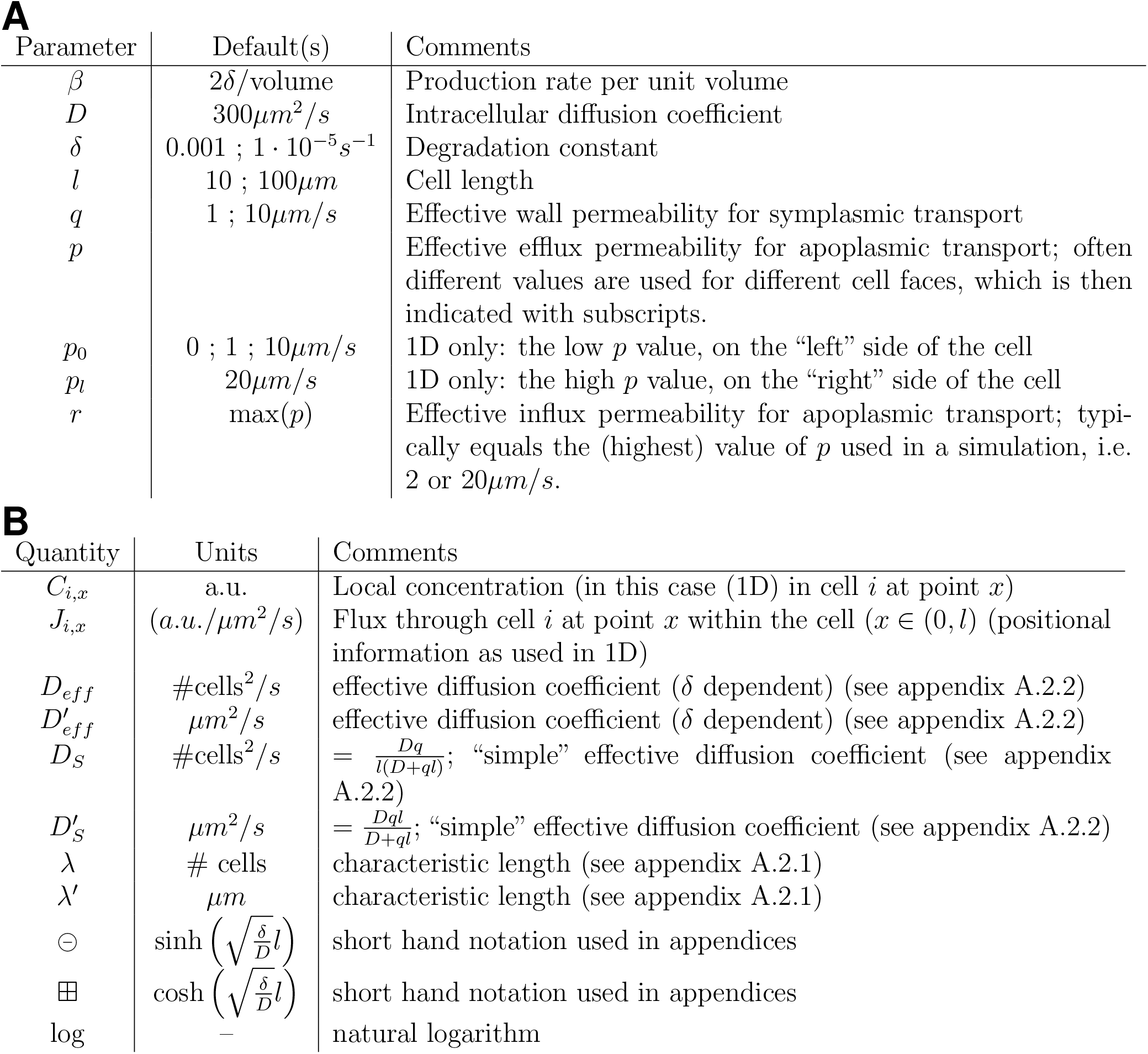
Overview of model parameters and mathematical symbols. **A**: parameters for symplasmic and/or apoplasmic transport. **B**: other quantities.

**A** Overview of mathematical symbols
| Parameter | Default(s) | Comments |
| --- | --- | --- |
| $\beta$ | $2\delta/\text{volume}$ | Production rate per unit volume |
| $D$ | $300\mu\text{m}^2/\text{s}$ | Intracellular diffusion coefficient |
| $\delta$ | $0.001 ; 1 \cdot 10^{-5}\text{s}^{-1}$ | Degradation constant |
| $l$ | $10 ; 100\mu\text{m}$ | Cell length |
| $q$ | $1 ; 10\mu\text{m}/\text{s}$ | Effective wall permeability for symplasmic transport |
| $p$ | | Effective efflux permeability for apoplasmic transport; often different values are used for different cell faces, which is then indicated with subscripts. |
| $p_0$ | $0 ; 1 ; 10\mu\text{m}/\text{s}$ | 1D only: the low $p$ value, on the “left” side of the cell |
| $p_l$ | $20\mu\text{m}/\text{s}$ | 1D only: the high $p$ value, on the “right” side of the cell |
| $r$ | $\max(p)$ | Effective influx permeability for apoplasmic transport; typically equals the (highest) value of $p$ used in a simulation, i.e. 2 or $20\mu\text{m}/\text{s}$. |

| Quantity | Units | Comments |
| --- | --- | --- |
| $C_{i,x}$ | a.u. | Local concentration (in this case 1D) in cell $i$ at point $x$ |
| $J_{i,x}$ | $(\text{a.u.}/\mu\text{m}^2/\text{s})$ | Flux through cell $i$ at point $x$ within the cell ($x \in (0, l)$) (positional information as used in 1D) |
| $D_{eff}$ | $\#\text{cells}^2/\text{s}$ | effective diffusion coefficient ($\delta$ dependent) (see appendix A.2.2) |
| $D'_{eff}$ | $\mu\text{m}^2/\text{s}$ | effective diffusion coefficient ($\delta$ dependent) (see appendix A.2.2) |
| $D_S$ | $\#\text{cells}^2/\text{s}$ | $= \frac{Dq}{l(D+ql)}$; “simple” effective diffusion coefficient (see appendix A.2.2) |
| $D'_S$ | $\mu\text{m}^2/\text{s}$ | $= \frac{Dql}{D+ql}$; “simple” effective diffusion coefficient (see appendix A.2.2) |
| $\lambda$ | $\# \text{ cells}$ | characteristic length (see appendix A.2.1) |
| $\lambda'$ | $\mu\text{m}$ | characteristic length (see appendix A.2.1) |
| $\ominus$ | $\sinh\left(\sqrt{\frac{\delta}{D}}l\right)$ | short hand notation used in appendices |
| $\boxplus$ | $\cosh\left(\sqrt{\frac{\delta}{D}}l\right)$ | short hand notation used in appendices |
| $\log$ | – | natural logarithm |

